# V1 interlaminar coherence decreases with interocular conflict

**DOI:** 10.1101/2025.10.04.680461

**Authors:** Brock M. Carlson, Blake A. Mitchell, Jacob A. Westerberg, Peter G. Poggi, Geoffrey Woodman, Alexander Maier

## Abstract

Resolving conflicting input from the two eyes is a fundamental challenge for the visual system. In the primary visual cortex (V1), such interocular conflict induces modest suppression of single neuron spiking, but the accompanying population-level dynamics remain poorly understood. Here we examined laminar multi-unit activity and interlaminar local field potential (LFP) coherence in macaque V1 during dichoptic stimulation and binocular rivalry flash suppression (BRFS). Using laminar microelectrode recordings, we found that interocular conflict reliably reduces interlaminar coherence, particularly between granular and infragranular layers, suggesting altered temporal coordination across the cortical column. Strikingly, during BRFS, coherence remained reduced even when firing rates were largely unchanged. Moreover, interlaminar coherence is higher for perceptually dominant BRFS stimuli, indicating that coherence across V1 layers covaries with perceptual outcome in the absence of significant firing-rate differences. These findings suggest that the temporal dynamics of V1 population activity are a more robust signal of interocular conflict than spike rate modulation.

**NEW & NOTEWORTHY:** These findings suggest that interocular conflict is processed across V1’s cortical layers through temporal coordination as well as modest rate changes of population activity. Interlaminar coherence therefore offers a complementary perspective on V1’s role during binocular rivalry, providing insight into population dynamics that may shape how visual signals are relayed to subsequent stages of visual processing.

## INTRODUCTION

Binocular rivalry is a spontaneous fluctuation in visual awareness that arises from interocular conflict, and it has therefore often been utilized to explore the neuronal basis of perception in both neurophysiology recordings (1–7) and neuroimaging (fMRI) (8–13). Interocular conflict leads to suppressed spiking activity in the primary visual cortex (V1), a mechanism known as dichoptic suppression (14–20). However, when subjects are under anesthesia or divert their attention, these modulations in V1 persist, suggesting that the associated response magnitude changes are not tied to perceptual outcome (21, 22). We aim to further characterize these circuit dynamics by examining the laminar activity in V1 during interocular conflict and binocular rivalry flash suppression (BRFS). We use two methods to investigate V1 laminar circuitry: multi-unit spiking activity (MUA) and interlaminar coherence.

Prior work has characterized spiking responses in V1 during both rivalry and BRFS using single electrodes (3, 11). However, due to technical limitations at the time, these neuronal responses were collected without locating them to specific cortical layers. As laminar computations are central to V1(23, 24), previous studies may not have been able to observe the full gamut of neuronal processes associated with interocular conflict. First, we utilized laminar electrodes to record MUA from V1 while macaques fixated on interocular conflict and BRFS. Second, we examined interlaminar coherence of the local field potential (LFP) as we were interested in whether the temporal structure of extracellular voltage fluctuations was informative about V1 processing conflict between the eyes. Previous studies on rivalry have shown that blood oxygen level dependent (BOLD) signals in V1 are more strongly coupled to local field potentials (LFPs) than to spiking activity (25, 26). As such, using a measure such as interlaminar coherence that evaluates the relationship between known causal components (spiking) and a reflection of the temporal structure of their population activity (via LFP) could provide a useful bridge in explaining results across methodologies (25, 27).

It has previously been proposed that conflict between the eyes could affect the coherence of V1’s output signals (28, 29), because synchronized neuronal populations firing can drive subsequent processing stages more effectively (30, 31). Hence, V1 coherence might serve as a mechanism to enhance processing of one eye’s information, without any concomitant change in average firing rate (19, 30, 32–35). Previous work in strabismic cats demonstrated that alternating states of binocular rivalry correlated with increased gamma-range synchrony but not with firing rate (34).

Certain aspects of neuronal population responses are invisible to the study of one neuron at a time, or even pairs of neurons at a time (35–42). Synchronization of oscillatory responses in cat area 17 provides insight into unit-to-unit response dynamics (34), but it remains unclear how this affects neurons at the scale of populations. Evaluating magnitude-squared coherence for V1 LFPs can serve as a heuristic for the laminar structure of coherence within single cortical columns (26, 43). We hypothesize that the temporal dynamics of V1 interlaminar interactions are disrupted during dichoptic (conflicting stimuli between the eyes) compared to dioptic (identical stimuli in each eye) visual stimulation. We found that during interocular conflict both spiking and interlaminar coherence decreased in V1. Likewise, we found that V1’s interlaminar coherence decreased whenever a column’s preferred stimulus was subject to BRFS inducing perceptual suppression – even though the average neuronal population activity remained largely unchanged. These new analyses of binocular rivalry responses from laminar electrodes in V1 identify modified interlaminar temporal dynamics during processing of interocular conflict.

## MATERIALS AND METHODS

### Animal Care

Neurophysiological recordings were obtained from two adult male macaques: monkey J, a rhesus macaque (Macaca mulatta), and monkey B, a bonnet macaque (Macaca radiata). Animal care and surgical procedures were approved by the Vanderbilt University Institutional Animal Care and Use Committee (IACUC) and conducted in accordance with National Institutes of Health and US Department of Agriculture (USDA) guidelines. Additional surgical and animal-care details are provided in the Supplemental Methods.

### Visual stimulation

#### Experimental setup

Visual stimuli were presented on a VIEWPixx display (VPixx Technologies; 120-Hz refresh rate, 1920 × 1200 pixels) through a custom Wheatstone-type stereoscope. The stereoscope used infrared-transmitting cold mirrors, permitting binocular stimulus presentation while eye position was monitored with an EyeLink II infrared eye tracker. Before experimental recordings, animals were trained to initiate fixation and maintain gaze within 0.5° of visual angle of a centrally presented fixation cue. The stereoscope was aligned separately for each recording session to match the visual positions seen through the two eyes (see Supplemental Methods).

Recording chambers provided access to the lower contralateral visual field representation of parafoveal V1. Receptive fields were in the left visual field for monkey “B” (right-hemisphere chamber) and in the right visual field for monkey “J” (left-hemisphere chamber). Based on the response properties of the neurons recorded from each acute electrode penetration, receptive-field eccentricities ranged from approximately 1.5° to 8° of visual angel. Most receptive fields were between 2° and 3° of eccentricity. Grating diameters scaled with eccentricity to match receptive-field location and size, ranging from 0.5° to 2.5° across experimental sessions. Receptive fields were mapped using localized noise patches and visual gratings, followed by drifting-grating measurements of orientation and eye preference.

Before beginning an experimental penetration, we required overlapping receptive fields and a common preferred orientation for more than 80% of the V1 recording sites along the probe, indicating that the contacts sampled a single cortical column. The orientation preferred by most sites was selected online as the preferred grating for the BRFS experiment; the orthogonal orientation served as the non-preferred grating. After acquisition, eye and orientation preferences were classified for individual multiunits using their monocular responses in the BRFS dataset. Thus, all stimulus comparisons were defined relative to each recording site’s measured preferences rather than assuming that every site shared the penetration’s predominant preference.

#### Experimental paradigm: Binocular rivalry flash suppression

The BRFS experiment used static sinusoidal gratings, generally at a spatial frequency of 1 cycle/degree and 50% Michelson contrast. When both eyes received a grating, the contrasts were matched. Visual stimulation and behavioral control were implemented using NIMH MonkeyLogic 2(44) and custom MATLAB scripts. The experiment contained 20 stimulus conditions: four simultaneous binocular conditions (dioptic or dichoptic), eight conditions with monocular adaptation followed by binocular stimulation, and eight monocular-alternation control conditions. Orientation and stimulated eye were counterbalanced in the condition definitions, and trial conditions were presented in pseudorandom order.

During BRFS, one eye viewed a grating for 800 ms; an orthogonal grating then appeared in the previously unstimulated eye for an additional 800 ms, while the original grating remained unchanged. There was no intervening blank interval. In the control condition of “monocular replay”, the original grating was instead removed at the transition, so that the visual stimulation physically alternated between the eyes. Trials were aborted when fixation was broken. Because aborted trials could yield unequal numbers of completed observations, experimental sessions were continued until at least 30 successfully completed trials had been obtained for each condition. Sessions that did not meet this minimum in all conditions were excluded from analysis.

### Neurophysiology

#### Recordings

Acute neurophysiological recordings were obtained using linear multielectrode arrays containing 32 contacts spaced 100 µm apart. Signals were acquired with a Blackrock Cerebus system at 30 kHz and subsequently processed to obtain local field potentials (LFPs) and multiunit activity (MUA; see Supplemental Methods).

We recorded 44 acute laminar penetrations from two macaques (12 in monkey B and 32 in monkey J). We determined the laminar position of each probe relative to V1 using established techniques (26, 43, 45–47). In particular, the bottom of the granular compartment (layer IVc) corresponded to the early stimulus-evoked current-source-density (CSD) sink (45, 46), supported by the transition between the alpha–beta and theta–gamma power spectral density (PSD) profiles (47) and the characteristic laminar coherence pattern. We also mapped receptive fields along the probe and evaluated whether the putative V1 sites had overlapping receptive fields and whether more than 80% shared the predominant orientation preference, consistent with recording within a single cortical column (14, 43).

Using these criteria, 31 of the 44 penetrations satisfied our initial laminar-alignment requirements. We then restricted the sample to penetrations containing at least five contacts in each of three operationally defined 0.5-mm laminar compartments: supragranular (layer II/II), granular (layer IV), and infragranular (layer V/VI). Four additional penetrations did not meet this depth-coverage criterion, leaving 27 penetrations for the final analyses (eight in monkey B and 19 in monkey J). These successive selection steps ensured reliable V1 alignment and balanced sampling of 15 contacts per penetration, yielding 405 MUA recording sites (135 per compartment).

#### Interlaminar coherence

Coherence analysis was used to examine the frequency-dependent LFP synchronization between pairs of electrode contacts within V1. The mscohere.m function in MATLAB was employed to assess magnitude-squared coherence. Coherence was evaluated over the final 512 ms of dichoptic stimulation to focus on sustained neuronal responses and minimize the influence of stimulus-onset transients. This approach is consistent with previous BRFS investigations that excluded early stimulus-evoked responses from their analyses (3, 7, 11, 22). The median coherence value across the frequency spectrum of 1-30Hz was obtained for each electrode pair on each trial. Median coherence values were then averaged across trials and recording penetrations.

To assess binocular subadditivity, we compared MUA responses to monocular stimulation of the preferred and non-preferred eyes with simultaneous dioptic stimulation of the same orientation. Monocular responses were evaluated during the initial 800-ms stimulus presentation, which was physically identical across four experimental conditions for each eye-orientation configuration. Monocular trials were randomly subsampled to match the number of dioptic trials within each recording penetration. Statistical comparisons were performed using paired t-tests across the sustained 512-ms response window (288–800 ms) and across eight 100-ms time bins, with Bonferroni correction for the latter comparisons. We additionally compared interlaminar coherence between preferred-eye monocular and dioptic stimulation during the same sustained response window. For the coherence analysis, stimulus preferences were determined from LFP responses, and the predominant preference across recording contacts was used to select the corresponding stimulus conditions for each penetration. Coherence was calculated using the same parameters as the primary analyses, with monocular trial counts matched to the available dioptic trials.

## RESULTS

### Laminar alignment

Laminar alignment was conducted on 44 acute electrode penetrations by determining the position of layer IVc using LFP, current source density (CSD), power spectral density (PSD), and MUA activity (46, 47). This alignment ensured accurate identification of cortical layers across penetrations. The relative laminar positions of each penetration were aligned to the bottom of the granular input layer. **Figure 1** shows an example laminar alignment for a single recording session. Traditionally, laminar alignment is accomplished with CSD (45, 46, 48) to identify the bottom of layer IVc initial stimulus-evoked current sink. The black arrow in **Figure 1** shows the channel that corresponds to the bottom of the CSD sink. However, utilizing laminar LFP power spectral density (PSD) can provide additional information.

**Figure 1:**
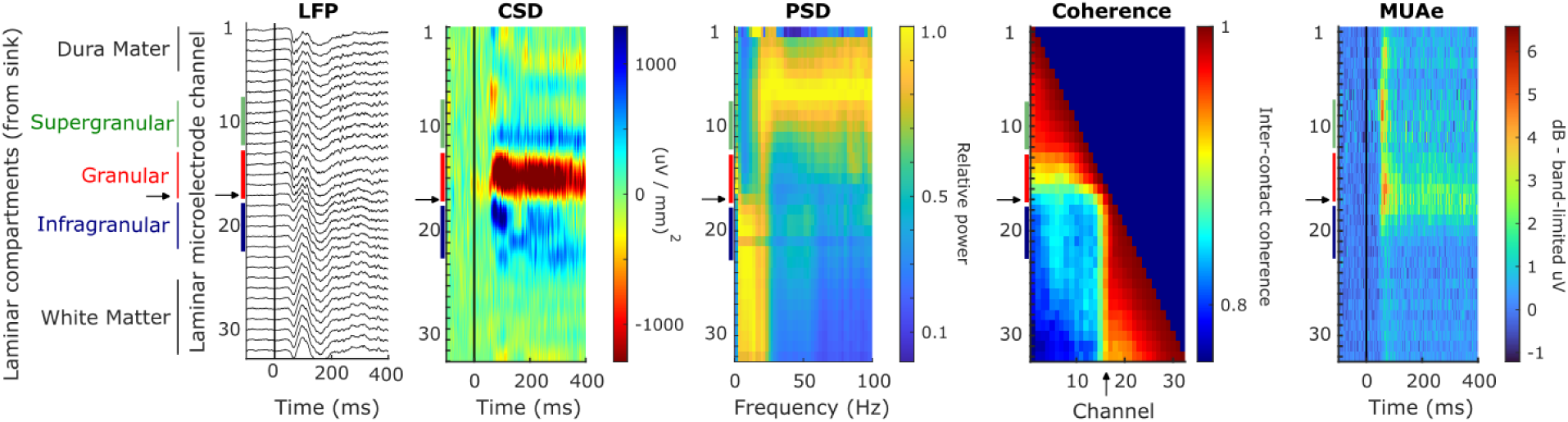
Laminar alignment. Example recording session. All 32 laminar microelectrode contacts are shown with complementary neurophysiological techniques: local field potential (LFP), current-source-density (CSD), LFP power spectral density (PSD), LFP coherence, and multi-unit spiking activity (MUA). The stimulus was a binocular grating of the column’s preferred orientation to maximally drive the response. The arrow at channel 17 shows where we determined that the transition between granular and infragranular tissue occurs. This determination was made primarily with CSD, PSD, and coherence analyses. From the bottom of the granular layer (the bottom of layer IVc), we established laminar compartments in units of 500μm or 5 electrode contact. The supragranular compartment (layer II/III) is indicated in green, the granular (layer IV) in red, and infragranular (layer V/VI) in navy. *Leftmost*: Stimulus-evoked laminar LFP response. *Second to left*: CSD response. A large and maintained initial current sink is observed in the granular layer (IVc) in the CSD, with a smaller transient sink in the supragranular layer. *Center*: The PSD shows the characteristic pattern of theta and gamma activity in the upper layers and alpha and beta activity in the deep layers. *Second to right*: LFP interlaminar coherence demonstrates the characteristic “step” that has previously been shown to co-locate with the granular-infragranular boundary. *Rightmost*: MUA confirms that stimulus-evoked spiking activity occurs throughout the entire length of the V1 column, and is most sustained within the granular layers, but does not have any reliable neurophysiological markers that correspond to laminar position.

Normalized theta and gamma power reliably peak in the upper layers and alpha and beta power peak in the deep layers (47). As a consequence, PSD can help identify deep penetrations within a fold of V2 or V3, which is something that CSD cannot do (47). We further verified our laminar alignment technique by utilizing magnitude-squared coherence (26). Paired coherence values for all electrode pairs at frequencies under 30Hz reveal a characteristic geometric pattern where coherence abruptly drops off at the layer IV/V boundary (26, 43). Multi-unit-activity (MUA) was defined by applying a high-frequency band-pass filter between 750-5000Hz to the broadband extracellular voltage signal, followed by half-wave-rectification, and then a bidirectional first-order low-pass filter to construct an “envelope” of spectral power (49, 50). MUA is considered to provide an aggregate measure of neuronal firing rates of neuronal populations less than 100 μm away from the electrode contact (50, 51). While MUA is useful for understanding the local output of neuronal responses, it does not exhibit a laminar morphology that can consistently be used for cross-session depth alignment in the same way as CSD, PSD, and coherence **(Figure 1).**

#### Effects of BRFS adaptation in V1

After laminar alignment of recording sessions, we examined the modulation of V1 neuronal firing rates that occur during BRFS. Our specific purpose was to examine how interocular conflict modulates responses across V1 laminar cortical columns. We use BRFS because it largely bypasses traditional rivalry’s initial mixed percept phase, producing almost immediate perceptual alternations (52–54). BRFS also leverages a brief period of monocular adaptation before presenting the dichoptic stimulus to generate a time-locked perceptual reversal, which is reliable enough (>90%) in both humans and primates that it has been previously used without subjective report (6, 7, 22, 29, 49, 54–59). Despite these advantages of BRFS, neural activity within the first ∼150 ms post-stimulus is considered pre-perceptual, reflecting automatic visual processes rather than perceptual processing (60, 61), since conscious perception lags the onset of visual presentations by at least this amount (62–64).

From 44 acute penetrations we identified 27 good recordings that spanned the full depth of V1 cortical columns with at least 5 electrode contacts in each compartment (N=8 in “B” and N = 19 in “J”), see Methods and SI. We verified that we were recording perpendicular to the cortex by observing overlapping receptive fields and shared orientation tuning along the probe.

We then determined how MUA responses vary across their most excitatory and least excitatory monocular stimulation conditions: the combination of their preferred and non-preferred eyes, as well as their preferred and orthogonal (non-preferred) orientations **(Figure 2A)**. We evaluated 27 penetrations, each with 15 electrode contacts. The grand average of our 405 V1 multi-units shows that population responses exhibited a robust preference for when their preferred eye and preferred orientation were monocularly stimulated **(Figure 2B),** consistent with prior work on ocular dominance (14, 65–67).

**Figure 2:**
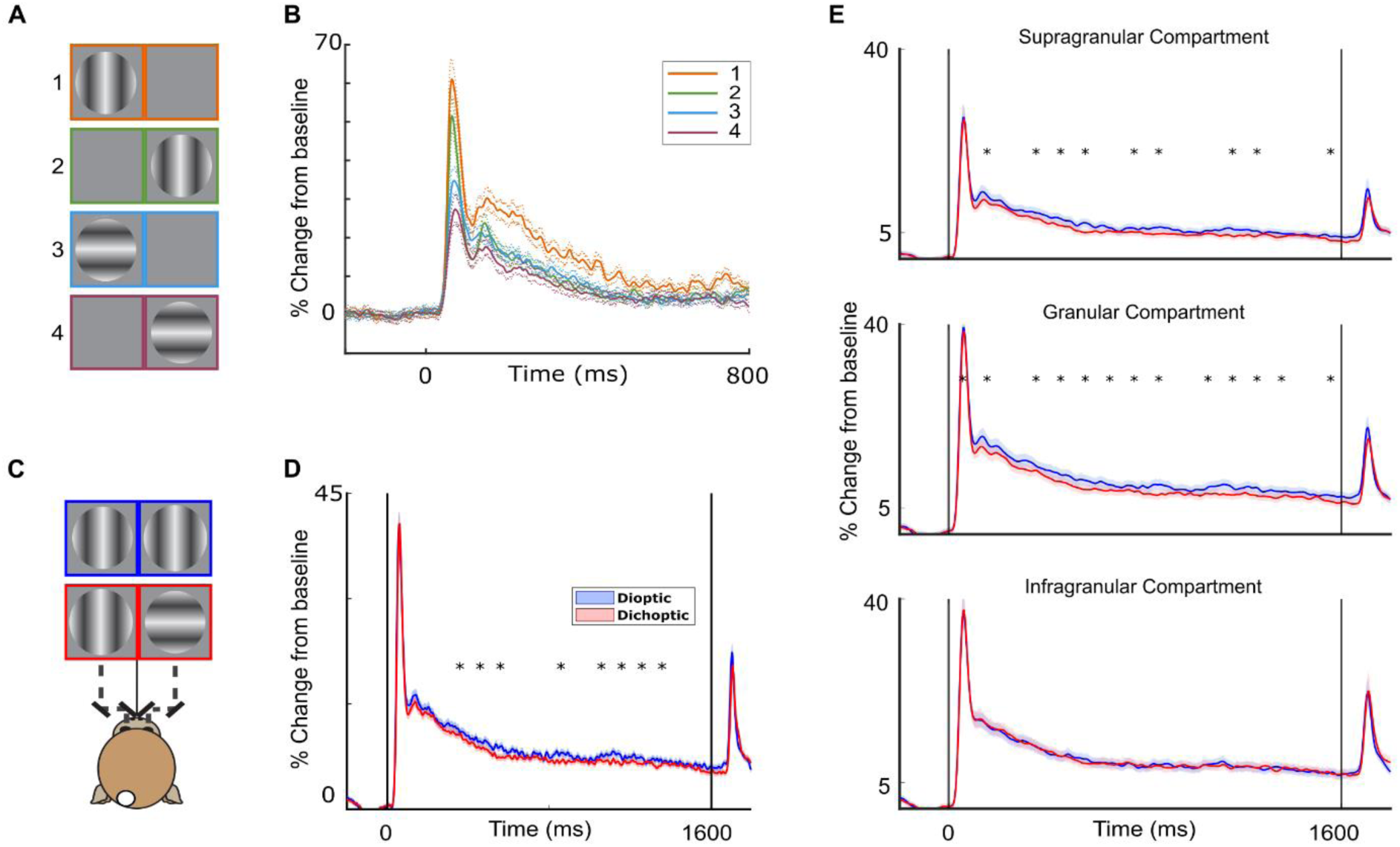
V1 population spiking responses to prolonged monocular, dioptic, and dichoptic stimulation. (A) Cartoon depiction of the monocular static gratings used to determine the ocular preference and orientation tuning. (B) Grand-average of 405 multi-units (both animals) evoked by monocular stimuli; neuronal activity is plotted as a mean percent change from baseline to normalize each unit’s activity across the population. Shaded regions on MUA plots represent standard error of the mean (SEM). Each multiunit has a preferred stimulus orientation and a preferred eye, as indicated by the orange line. (C) Cartoon depicting the mirror stereoscope used to present stimuli to the two awake and behaving macaques. The monitor was divided so that each eye could only see one half of the screen. The cartoon depicts an example dioptic stimulus (blue) and a dichoptic (red) stimulus. (D) Interocular conflict with dichoptic stimuli (red) produces a small yet reliable amount of dichoptic suppression with respect to dioptic stimuli (blue). Asterisks indicate 100ms time-bins with a statistically significant difference between the dioptic and dichoptic conditions (one-tailed paired t-tests, Bonferroni-corrected for 16 comparisons; α = 0.05). (E) V1 laminar recordings were performed with laminar microelectrodes at 100μm spacing. Multi-unit responses of contacts within each the supragranular, granular, and infragranular compartments were averaged.

Interocular suppression is a small yet reliable reduction in neural response that occurs in almost all V1 cells, decreasing their response rate when conflict arises between the eyes (14–16, 18, 49, 68–71). We replicated these findings in our dataset by measuring the percent change from baseline over the last 500 ms of the sustained neural response for each unit. This revealed a slight decrease in activity for the dichoptic condition compared to the dioptic condition (**Figure 2D**). Specifically, the dioptic stimulus produced an average 7.42% change from baseline (SD = 6.83%), whereas the dichoptic condition produced a 6.97% change (SD = 6.85%), resulting in a mean difference of 0.45% (SE = 0.19%), *t*(404) = 2.26, *p* = 0.012. This finding confirms that dichoptic suppression has a small effect size, requiring experiments to be adequately statistically powered to observe it (14, 49).

To study the time course of these neural responses, we divided the response period into 16 contiguous 100-ms bins, ranging from 0 to 1600ms. For each bin, we computed the mean firing rate (in percent change from baseline) of each MUA and performed a one-tailed paired t-test comparing the dioptic and dichoptic conditions. To correct for multiple comparisons across the 16 bins, we applied a Bonferroni correction. We observed modest but consistent dichoptic suppression, beginning approximately 200 ms after stimulus onset and persisting through the sustained response.

We next divided our 405 multi-units into laminar compartments, resulting in 135 recording sites in the supragranular, granular, and infragranular compartments, respectively. The strongest response differences to orthogonal eye and orientation features occur across distinct cortical columns, and are theorized to be most pronounced in the superficial layers II/III (72–76). Therefore, we expected that dichoptic suppression might occur most predominantly within the upper layers. To evaluate this hypothesis, we repeated the moving time-window analysis (as in **Figure 2D**) on each laminar compartment’s neuronal sub-population. We observed statistically significant layer-specific dichoptic suppression in the supragranular and granular compartments (**Figure 2E**). As hypothesized, the deepest layer of cortex did not show significant dichoptic suppression.

#### Inter-Laminar Coherence Reflects Interocular Conflict

Given the distinct laminar profile of dichoptic suppression, we wanted to closely examine how interocular conflict alters information flow *between* cortical layers. To address this question, we calculated inter-laminar LFP coherence to provide insight into the temporal coordination of neuronal activity along the V1 cortical column. For example, it has previously been hypothesized that V1 may engage in the processing of binocular rivalry through coordinating the temporal dynamics of population firing rather than signaling that information by the spike rate alone (11, 28, 77). In support of this hypothesis, it has been shown that neural synchrony decreases with interocular conflict in anesthetized strabismic cats (34). We expanded on this work by evaluating magnitude-squared coherence between each V1 laminar electrode contact. We specifically hypothesized that interlaminar coherence is higher for dioptic than for dichoptic stimulation, following the associated changes in average firing rate. Indeed, laminar coherence proved higher during dioptic stimulation than during dichoptic stimulation **(Figure 3)**.

**Figure 3:**
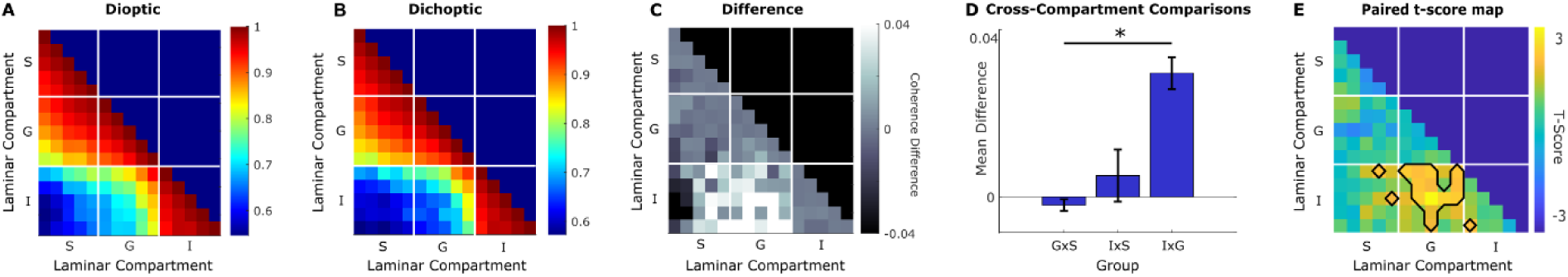
Laminar LFP coherence reflects interocular conflict. Interlaminar LFP coherence is calculated for the last 512 ms of the visual response on each trial. Values were averaged across trials within each condition and across 27 penetrations. Magnitude-squared coherence shows the degree to which the LFP signal from one channel relates to the LFP signal from another channel along the electrode. Based on prior work (26), we expected a clear structural pattern of V1 columns, where supragranular and granular layers show coherence within themselves and the infragranular layer is mostly coherent only with itself. Dioptic (A) and dichoptic (B) conditions look similar in their overall structure, but their difference (C) shows that coherence enhancement was greater during dioptic compared to dichoptic stimulation, and to a greater extent between the infragranular and the granular layers. The labels S, G, and I refer to supragranular, granular, and infragranular compartments. (D) ANOVA on coherence differences between cross-compartment comparisons. The asterisk indicates a significant effect of the interlaminar compartment on the coherence differences between conditions (one-way ANOVA, *F*(2,72) = 14.83, p < 0.001). (E) The infragranular to granular coherence difference is clear in the map of *t* scores for the paired coherence difference. Outlined electrode channel pairs indicate a statistically significant difference between dioptic and dichoptic coherence (paired t-test, α = 0.05).

Interlaminar coherence has a consistent pattern with respect to cortical depth that arises independent of behavioral state or cortical stimulation (26, 43). This finding holds, as the *overall* pattern of coherence remains similar between dichoptic and dioptic stimulation **(Figure 3A and 3B)**. It was only in the *contrast* of responses to the two types of binocular stimulation that interocular conflict decreased interlaminar coherence **(Figure 3C).** To quantify this difference, we performed a one-way ANOVA over the coherence values between each laminar compartment. Because each compartment contained five electrodes, each pairwise comparison yielded 25 coherence values (5 × 5) for each of the three interlaminar comparisons. The ANOVA demonstrated a significant main effect of compartment (F(2, 72) = 14.831, p = 4.039×10−6) (**Table 1**). Notably, the largest decrease in coherence occurred as soon as the boundary below layer IVc was crossed (**Figure 3D and 3E**), indicating that conflict-related changes are especially pronounced when comparing the granular layer to the infragranular layer. While one might expect the greatest difference to arise between the supragranular and infragranular layers due to their relative spatial distance. These results highlight the importance of the layer IV/V boundary for interlaminar coherence.

**Table 1:** ANOVA for interlaminar coherence decrease with interocular conflict.

|  | **SS** | **Df** | **MS** | **F** | **p** |
|---|---|---|---|---|---|
| Groups | 0.01368 | 2 | 0.00684 | 14.831 | 4.0387e-06 |
| Error | 0.03320 | 72 | 0.00046 |  |  |
| Total | **0.04688** | **74** |  |  |  |

#### Binocular Rivalry Flash Suppression

Dichoptic stimulation evokes the perceptual phenomenon of binocular rivalry (BR). BR consists of a largely random succession of conflicting stimuli. However, probing an animal’s visual perception during binocular rivalry poses significant methodological challenges (78). To address these concerns, we utilized binocular rivalry flash suppression (BRFS), which is one of the most frequently used variants of binocular rivalry for animal experiments (79). Unlike conventional BR, which results in stochastic visual changes (80–82), the phenomenology of BRFS is predictable (52, 83). In BRFS, a period of monocular adaptation to one eye is followed by the immediate onset (flash) of binocular dichoptic presentation (54). When the dichoptic flash occurs, perception reliably elevates the unadapted stimulus presented to the unadapted eye to perceptual dominance (11, 54). Most previous binocular rivalry studies in macaques have relied on BRFS, establishing psychophysical support that macaques experience BRFS similar to humans (3, 7, 15, 16, 18, 22, 28). We first replicated prior results **(Figure 4A).** Then, we expanded on past work by analyzing neuronal firing across the entirety of the V1 cortical column, as shown in **Figure 4B**.

**Figure 4:**
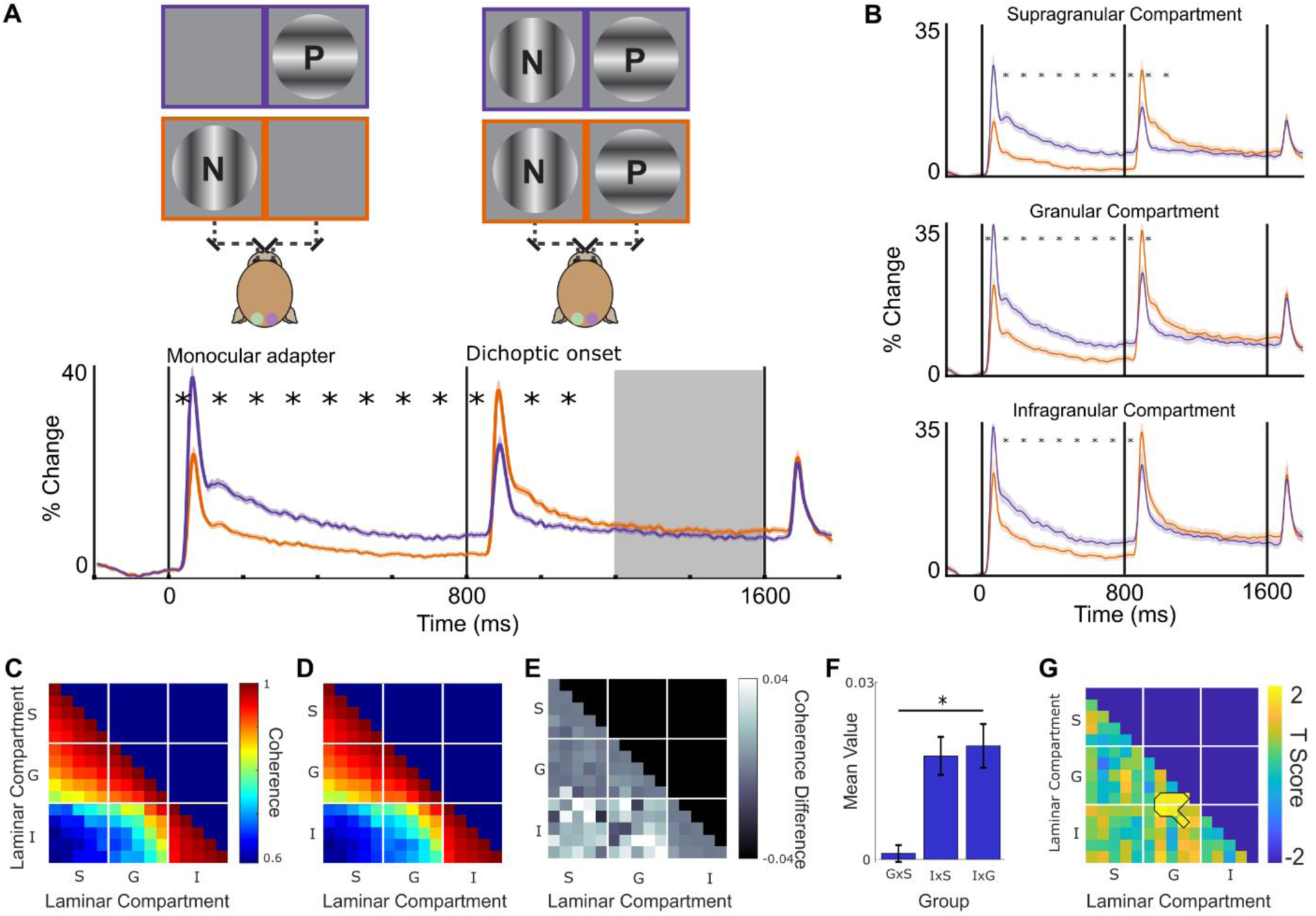
Binocular rivalry flash suppression (BRFS). (A) Cartoon showing BRFS stimuli and plot showing the associated MUA responses. Left and right grey squares in the cartoon indicate halves of the monitor split by the stereoscope. BRFS consisted of monocularly adapting one eye with either the column’s preferred (purple) or null (orange) stimulus and then flashing a dichoptic stimulus (purple and orange) at 800ms. At the moment of the dichoptic flash the visual stimulation is identical between the purple and the orange condition. Solid lines represent the mean response of 405 multi-units. Shaded areas represent SEM. Asterisks indicate 100ms time-bins with a statistically significant difference between the two BRFS configurations (one-tailed paired t-tests, Bonferroni-corrected for 16 comparisons; α = 0.05). (B) The V1 laminar compartments appear largely homogeneous in their spiking response 200ms after the dichoptic onset. (C) Interlaminar coherence for the preferred stimulus onset during the last 512ms of BRFS (orange line). (D) Same as in C, but for the non-preferred stimulus onset (purple line). (E) Coherence difference between C and D. (F) Cross-compartment coherence distributions demonstrate that there is significantly higher interlaminar coherence when the preferred dichoptic stimulus is presented second than when the null dichoptic stimulus is presented second. ANOVA on coherence differences between compartments. The asterisk indicates a significant effect of interlaminar compartment on the coherence differences between conditions (one-way ANOVA, F(2,72) = 11.63, p < 0.001). (G) Paired t-score map shows that significant differences lie along the granular and infragranular boundary. Outlined electrode channel pairs indicate a statistically significant difference between BRFS when the preferred stimulus is perceived vs when the null stimulus is perceived (paired t-test, α = 0.05).

The 800 ms of monocular adaptation is crucial for our paradigm because it ensures a reliable perceptual effect of BRFS, whereas periods shorter than 500 ms can occasionally yield mixed percepts (54).

Adaptation has a well-documented suppressive impact on V1 neuronal activity (43, 57, 84–92). The influence of adaptation in BRFS is evident in the distinct neural *transients* elicited by the dichoptic onset. Despite identical retinal stimulation between BRFS conditions at 800ms, the strength of the dichoptic onset’s evoked response depends on the history of monocular adaptation (**Figure 4A)**. This effect of adaptation is short-lived, however, and subsides around ∼200ms, at which point perception of the visual stimulus starts to arise. At that point, the two responses become largely equal, reflecting the physical similarity of these stimulus conditions (i.e., the same stimuli presented to the same eyes).

We performed a moving time-window analysis to pinpoint the duration of the initial transient difference between the purple and orange conditions in both the grand average response **(Figure 4A)** and the laminar compartments **(Figure 4B).** As before, 16 100-ms time bins were analyzed with a t-test, and a Bonferroni correction was applied. As shown in **Figures 4A** and **4B**, the monocular adaptation period of BRFS evoked a significant difference in neuronal response. This significant difference persists through the transient onset of binocular stimulation until approximately 250 ms following dichoptic stimulus onset, after which the two BRFS conditions converged to similar sustained firing rates.

We then evaluated interlaminar coherence during the sustained firing window (last 512 ms of the stimulus) and observed a significant difference in interlaminar coherence between the BRFS conditions (**Figure 4C–F)**. Specifically, when the preferred stimulus was presented second, and thus assumed to dominate perception (orange traces in **Figure 4A**, coherence in **Figure 4C**), there was markedly higher interlaminar coherence between the deep and upper layers than when the non-preferred stimulus was presented second (purple traces in **Figure 4A**, coherence in **Figure 4D**). The coherence difference between BRFS conditions is evident in **Figure 4E**. This effect occurred during a period in which there was no statistically significant difference in the neuronal firing rates.

We performed a one-way ANOVA on the coherence values between laminar compartments. As summarized in **Table 2** (SS = 0.00506, MS = 0.00253, F(2,72) = 11.63, p < 0.001), the main effect of BRFS was significant. **Figure 4E and Figure 4F** demonstrate that both the infragranular-to-supragranular and infragranular-to-granular comparisons exhibit higher coherence when the preferred stimulus is presented second – and ostensibly perceived -compared to the null stimulus. **Figure 4G** illustrates precise laminar locations where a paired t-test reveals significant differences between the BRFS conditions (p < .05). These differences, similar to the dioptic and dichoptic comparison, are located on the granular to infragranular boundary.

**Table 2:** ANOVA for interlaminar coherence decreases with BRFS.

|  | **SS** | **Df** | **MS** | **F** | **p** |
|---|---|---|---|---|---|
| Groups | 0.00506 | 2 | 0.00253 | 11.63363 | 4.1879e-05 |
| Error | 0.01567 | 72 | 0.00022 |  |  |
| Total | 0.02073 | 74 |  |  |  |

To further characterize the frequency dependence of these effects, we repeated the BRFS coherence analysis separately for the alpha, beta, and gamma frequency bands (**Supplemental Figure 2**). The alpha- and beta-band analyses revealed patterns of interlaminar coherence differences similar to those observed in the primary 1–30 Hz analysis. The gamma-band analysis also revealed a significant effect of laminar compartment, although the coherence differences were in the opposite direction from those observed at lower frequencies. We also evaluated interlaminar coherence during the first 512 ms following the dichoptic onset (**Supplemental Figure 3**). However, this early window includes the large stimulus-evoked transient and is therefore more susceptible to nonstationarity in the LFP signal. Consequently, we retained the final 512 ms of stimulation as the primary analysis window, consistent with previous investigations of binocular rivalry and BRFS (3, 7, 11, 22).

To determine whether the coherence differences observed during BRFS could be attributed to the abrupt stimulus transition, we examined a monocular replay control condition in which stimuli physically alternated between the eyes without simultaneous dichoptic stimulation (**Supplemental Figure 4**). Unlike the sustained MUA responses during BRFS, monocular replay produced substantial differences in MUA between the preferred- and null-stimulus sequences (**Supplemental Figure 4A**). Interlaminar coherence also differed between monocular replay conditions, although the laminar distribution of these differences was visually distinct from that observed during BRFS (**Supplemental Figure 4B**). Specifically, the granular-to-infragranular had much less interlaminar coherence on average during monocular replay than during BRFS. These results demonstrate that interlaminar coherence can vary during physical stimulus alternation, while also indicating that the coherence changes observed during monocular replay do not simply reproduce the pattern found during BRFS.

#### Power Spectral Density

Thus far we have endeavored to demonstrate that the nature of interlaminar communication changes during dichoptic processing. Differences in interlaminar coherence may arise from changes in neuronal population dynamics, but they can also be influenced by differences in LFP power (33, 93). To evaluate whether changes in the relative distribution of LFP power could contribute to our observed coherence differences, we examined the laminar power spectral density (PSD) across stimulation conditions (**Figure 5**). All conditions exhibited the characteristic spectrolaminar pattern, with alpha- and beta-band power strongest in the infragranular layers and gamma-band power strongest in the granular and supragranular layers (47). The comparison of dioptic and dichoptic stimulation revealed relatively small differences in laminar PSD, including changes within the alpha-beta frequency range (**Figure 5C**). During BRFS, however, the relative laminar distribution of LFP power was nearly identical between conditions (**Figure 5F**), despite the differences in interlaminar coherence reported in **Figure 4**.

**Figure 5:**
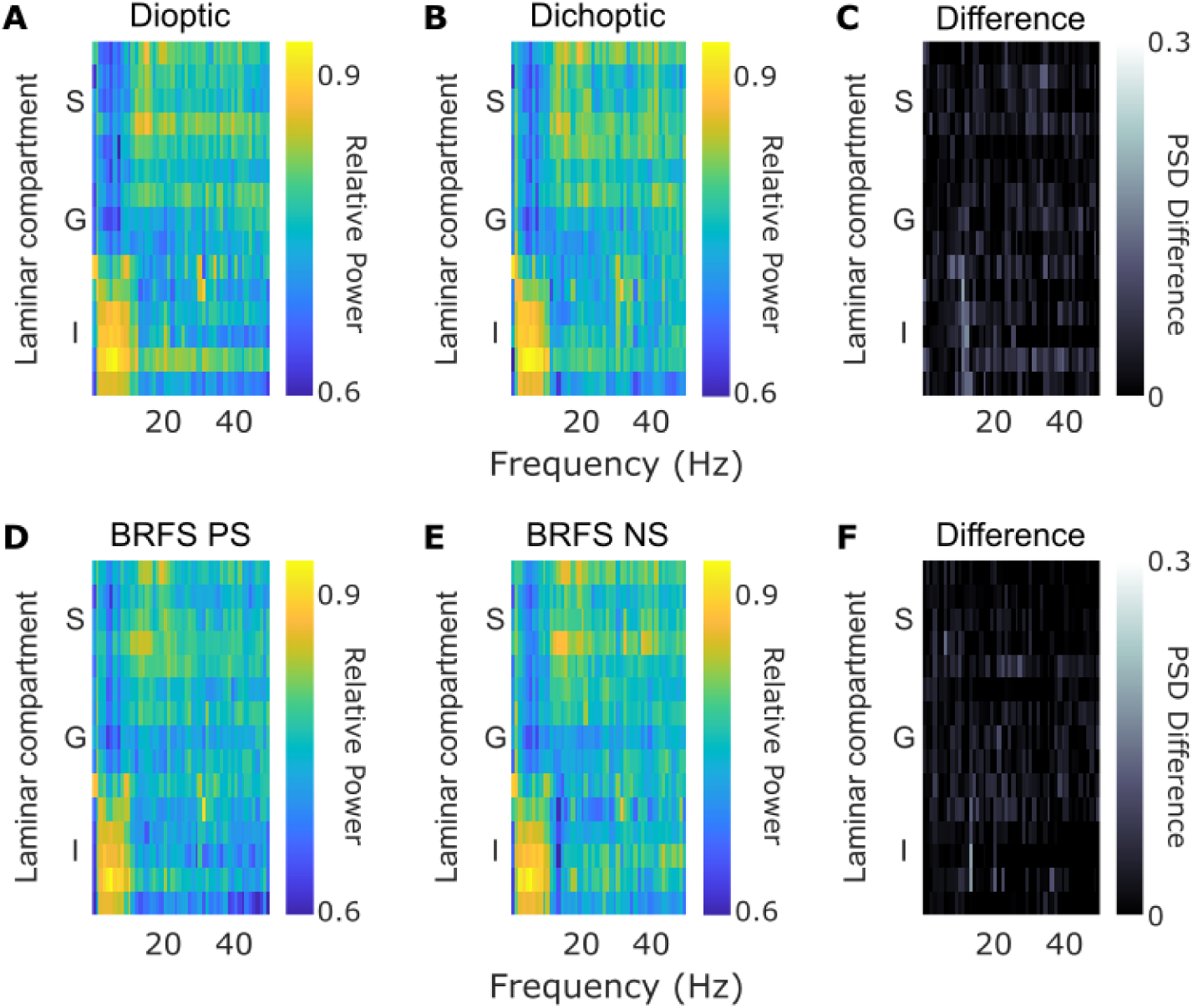
Laminar PSD is largely identical across conditions. Laminar power spectra were calculated for the last 512ms of the visual response on each trial. PSD was averaged across trials and then normalized across channels. For each frequency, the channel with the maximum power was assigned a value of 1, and all other channels were scaled relative to that maximum. Averages for each condition across 27 penetrations are shown. Dioptic (A) and dichoptic (B) conditions look similar in their overall structure, and their difference (C) shows very little change in the laminar PSD between the two conditions. Similarly, (D) shows the laminar PSD for the last 512ms of BRFS after the preferred stimulus is flashed, (E) shows the opposite BRFS condition where the null stimulus is flashed, and their difference (F) shows very little change in laminar PSD between the two conditions.

#### Binocular normalization

Finally, we evaluated whether binocular responses could be explained by the linear summation of their monocular components (**Supplemental Figure 5A**). For dioptic stimulation of the preferred orientation, the observed MUA response was substantially smaller than the sum of the responses to monocular stimulation of each eye (13.46% vs. 9.86% change from baseline; paired t(404) = 8.07, p < 0.001). This subadditivity was significant across all eight 100-ms time bins (one-tailed paired t-tests, Bonferroni-corrected for eight comparisons; α = 0.05). Furthermore, the binocular response closely resembled, yet was significantly greater, than the response to monocular stimulation of the preferred eye, consistent with previous reports of binocular normalization (66, 67, 94, 95). A test was done on the average MUA response over the sustained 512-ms window (9.86% vs. 8.11% change from baseline; paired two-tailed t(404) = −6.44, p < 0.001). This difference between dioptic and monocular preferred responses was also significant in seven of eight 100-ms time bins after Bonferroni correction.

We next compared interlaminar coherence during preferred-eye monocular and dioptic stimulation using the same sustained 512-ms analysis window (**Supplemental Figure 5B**). Coherence differences between these conditions varied significantly across interlaminar compartment comparisons (one-way ANOVA, F(2, 72) = 3.64, p = 0.031). Thus, as in the dioptic–dichoptic comparison (**Figures 2 and 3**), coherence differences between monocular and dioptic stimulation were accompanied by significant differences in MUA. In contrast, the BRFS response demonstrated interlaminar coherence differences despite the absence of significant firing-rate differences (**Figure 4)**.

## DISCUSSION

### Reduced laminar coherence reflects interocular conflict

Our sensory systems frequently receive conflicting information, and yet we maintain a coherent, unitary perceptual experience of the external world (96). Therefore, it is of great interest to study how the brain resolves conflicting inputs. Psychophysically, it is readily feasible to induce conflict between the eyes (97), and interocular conflict is an essential element of viewing natural scenes (98). As such, one of the most well-documented neuronal binocular mechanisms in V1 is dichoptic suppression, where conflict between the eyes results in a decrease in neuronal firing (15, 16, 18). Observed in cats (15–18, 99), macaques (14, 49), and humans (12, 21), interocular conflict reliably induces a small decrease in V1 neuronal population activity. It should be noted that the effect size of this neuronal suppression is comparatively small, yet reliable (49).

Previously, the onset of dichoptic suppression has been reported to start 250 ms after stimulus onset, suggesting that the neuronal suppression could be caused by feedback onto V1 (14). We observe a similar time course in our grand average of 405 V1 multi-units, replicating previous reports (15, 49). We expand on these findings by evaluating V1 spiking responses with respect to their laminar depth. In doing so, we observed heterogeneity in the laminar response profile of multi-unit activity to interocular conflict. Evaluated in sliding time-windows, corrected for multiple comparisons, population-based dichoptic suppression is exclusively observed among the supragranular and granular cell populations. In contrast, the infragranular units did not show significant dichoptic suppression on average. Dichoptic suppression, which occurred exclusively in the upper (feedforward-associated) layers, suggests that V1 columnar populations signal the presence of interocular conflict to the rest of the visual system via spiking activity in a feedforward fashion.

We also observed a decrease in interlaminar coherence during interocular conflict. Past work has highlighted a consistent pattern of interlaminar coherence across various stimulus types, including light flashes, static gratings, and blank screens (26). We expand upon this work by looking at coherence differences between stimulation paradigms. We demonstrate that interocular conflict can alter the magnitude of coherence within the V1 cortical column, particularly between the granular and infragranular layers. However, these coherence differences between dioptic and dichoptic stimulation were also accompanied by modest but significant differences in average MUA responses. Changes in interlaminar coherence under these circumstances are perhaps unsurprising, as changes in neuronal population activity may also alter the temporal relationships between cortical layers. Indeed, our monocular replay control demonstrates that substantial differences in physical stimulation can produce differences in both MUA and interlaminar coherence (**Supplemental Figure 4**). While these observations demonstrate that visual stimulation modulates multiple aspects of V1 population activity, they do not establish whether coherence provides information beyond that reflected in average firing rates. This distinction becomes particularly important when examining the sustained neuronal responses during BRFS.

### Interlaminar coherence decreases with BRFS suppression, while spiking activity remains largely unchanged

During BRFS, the onset of dichoptic stimulation following monocular adaptation evokes a transient difference in spiking activity within V1. Spiking is higher for columns whose preferred stimulus is presented second than for those adapted to the preferred stimulus. This difference is robust during the first ∼250 ms of the dichoptic presentation and can be most parsimoniously attributed to well-established mechanisms of neuronal adaptation (43, 49, 84, 87, 90). Adaptation reduces excitability in neurons exposed to sustained input, leading to a dampened response upon subsequent stimulation with the same features (86, 88, 92, 100).

Behavioral and neurophysiological studies estimate that conscious perception of a stimulus does not emerge until ∼250 ms after stimulus onset (14, 60, 61). It follows that this early response window likely falls outside the timeframe associated with visual awareness. Our goal in this manuscript is not to identify the neural correlates of consciousness (101–103) or to revisit previous debates about whether V1 plays a perception-determining role during rivalry (1, 21, 104, 105). Instead, we interpret the MUA differentiation during the first 250ms of BRFS’ dichoptic flash as reflecting pre-perceptual, reflex-like circuit dynamics. This interpretation is further supported by prior studies showing that rivalry induced neural responses in V1 can persist under conditions of diverted attention or anesthesia (21, 22).

Prior studies investigating perceptual suppression in BRFS and conventional binocular rivalry have limited their analysis to the sustained response window that follows the initial response transient (3, 7, 11, 22). This window also provides a relatively stationary signal for Fourier-based coherence analyses. Consistent with this approach, we examined the final 512 ms of the dichoptic stimulation period. During this interval, in agreement with previous work, average MUA responses were not significantly different between BRFS conditions (**Figure 4A–B**), and the relative laminar distribution of LFP power was nearly identical (**Figure 5D–F**). Nevertheless, interlaminar coherence remained significantly modulated, particularly between the infragranular and upper cortical layers (**Figure 4C–G**).

The supplemental analyses provide further context for interpreting this dissociation. Frequency-specific analyses revealed similar coherence modulation in the alpha and beta bands, although the direction of the coherence differences reversed in the gamma band (**Supplemental Figure 2**). The frequency dependence of these coherence effects is consistent with the established spectrolaminar organization of V1, in which alpha-beta power is strongest in the infragranular layers, whereas gamma power predominates in the superficial layers (47). This organization is also evident in the PSD observed in our recordings (**Figures 1 and 5**). Given that the largest coherence differences involve the infragranular compartment, the different patterns observed at lower and higher frequencies may reflect distinct frequency-dependent interactions across cortical layers. Coherence differences were also observed during the initial stimulus-onset transient (**Supplemental Figure 3**). The monocular replay control demonstrated that physical alternation between preferred and null stimuli induced substantial differences in both MUA and interlaminar coherence (**Supplemental Figure 4**). Similarly, the subadditivity analysis demonstrated that the binocular MUA response was substantially smaller than the linear sum of its monocular components (**Supplemental Figure 5A**), consistent with previous accounts of binocular integration and normalization (66, 67, 106, 107). An additional comparison of preferred-eye monocular and dioptic stimulation revealed modest, compartment-dependent coherence differences accompanied by significant changes in MUA (**Supplemental Figure 5B**). These observations suggest that binocular integration can influence both the magnitude and temporal relationships of neuronal population activity. However, because the monocular and dioptic conditions also differed in average MUA, this control does not establish whether normalization can independently modulate interlaminar coherence.

Together, these analyses demonstrate that changes in visual stimulation can influence both MUA and interlaminar coherence. Such concurrent changes are not unexpected, as differences in neuronal population activity may also alter the temporal relationships between cortical layers. The more striking observation arises during sustained BRFS, when interlaminar coherence continues to differentiate conditions despite the absence of significant differences in average neuronal firing rates.

This dissociation is the central finding of our study. Unlike the dioptic/dichoptic comparison, in which changes in coherence accompany changes in average firing rates, the sustained BRFS response reveals differences in coherence without corresponding significant differences in MUA. These findings suggest that the temporal coordination of V1 population activity reflects aspects of visual processing that cannot be characterized by average firing rates alone. Furthermore, traditional rate coding cannot fully account for the rivalry-induced neural dynamics observed in V1 (108).

The potential for interlaminar coherence to indicate a perceptually relevant signal thus remains an intriguing possibility (19, 26, 29, 33, 34, 109–111). Because the BRFS conditions shared identical dichoptic stimulation but differed in their adaptation history and expected perceptual dominance, the observed coherence differences may reflect V1 microcircuit activity relevant to perceptual suppression rather than merely changes in neuronal response magnitude. Interlaminar coherence therefore provides a complementary measure of V1 population dynamics, revealing aspects of visual processing that are not apparent from average population firing rates.

### LFP Coherence as Putative Reflection of Population Activity

The causal efficacy of the LFP is a matter of contention. Studies on ephaptic coupling have shown the potential for causal effects of LFP changes on modulating neuronal spike timing (rather than spike rate) (112, 113). However, the field strength required for this to occur has only been reported for rare instances *in vivo*, such as during epileptic seizures or in non-cortical structures (114–116). It thus seems unlikely (though it remains an open question) whether LFP can causally affect cortical spiking. However, even if assuming that LFP *magnitude* can impact spiking directly, it is less clear how LFP *coherence* might have direct causal impact on spiking activity (but see: (117)). LFP coherence might be more plausibly regarded as a (epiphenomenal) reflection of population-based dynamics (e.g., of synchronized synaptic processing) that could go unnoticed using limited-resolution techniques. The current trend of increasing electrode contact counts, from the double digits observed in this study to triple digits and beyond, coupled with high-density current source density (CSD) analysis, may enable a more direct investigation of the underlying synaptic processes.

### Population Signals in V1 During Interocular Conflict

Our findings demonstrate that interlaminar coherence can reveal changes in V1 population activity that are not apparent from average neuronal firing rates. Whereas conventional measures of spiking activity primarily characterize the magnitude of neuronal responses, coherence provides information about the temporal relationships between neuronal populations. Furthermore, interlaminar coherence demonstrates characteristic temporal interactions across the cortical columnar microcircuit. During BRFS, these relationships were altered despite the absence of significant differences in sustained MUA responses. Synaptic changes that remain subthreshold may be reflected in coherence across cortical layers. While firing-rate modulations are a well-established mechanism for sensory coding, these results indicate that the processing of interocular conflict involves changes in the temporal coordination of neuronal populations in addition to conventional rate-based responses.

## DATA AVAILABILITY

Source data for this study are openly available at 10.5281/zenodo.18602727. Analysis repository is available at https://github.com/BrockMCarlson/bmcBRFSanalysis.

## SUPPLEMENTAL MATERIAL

### Methods

#### a. Animal Care

Animal care procedures adhered to strict ethical guidelines and were approved by the Institutional Animal Care and Use Committee (IACUC) at Vanderbilt University, in accordance with National Institutes of Health regulations and the Association for the Assessment and Accreditation of Laboratory Animal Care (AAALAC). Two adult male macaques participated: monkey J (Macaca mulatta) and monkey B (Macaca radiata). Animals underwent sterile surgical procedures for implantation of head holders and recording chambers constructed from MRI-compatible materials. Isoflurane anesthesia (1.5–2.0%) was maintained during surgery, and physiological variables were continuously monitored. Recording chambers were positioned above the occipital ridge and lateral to the midline venous sinus to provide access to parafoveal V1. The chamber was implanted over the right hemisphere in monkey B and over the left hemisphere in monkey J. Chambers and head holders were secured with transcranial ceramic screws (Thomas Recording, Gießen, Germany) and dental acrylic (Lang Dental Manufacturing, Wheeling, IL). Postoperative care included analgesics, antibiotics, and veterinary monitoring. All recording penetrations were acute.

#### b. Experimental Paradigm

##### visual display, stereoscope, and fixation

Visual stimuli were shown on a VIEWPixx display (VPixx Technologies; 22.5-inch screen, 1920 × 1200 pixels, 120-Hz refresh rate). Images for the two eyes were presented on separate halves of the monitor, divided by a nonreflective septum, and viewed through a custom Wheatstone-type mirror stereoscope. Infrared-transmitting cold mirrors allowed the visual display to be reflected to each eye while infrared images of both eyes remained available to an EyeLink II eye tracker. Matching peripheral contours supported binocular fusion. At the start of each session, the mirrors were adjusted using fixation targets shown at corresponding locations to either eye, and eye-position measurements were used to confirm alignment. Animals were trained to initiate fixation on a central cue and maintain gaze within 0.5° of visual angle. Trials in which fixation was not acquired or was subsequently broken were aborted.

##### Receptive-field mapping and stimulus selection

The recording chambers targeted the dorsal, parafoveal representation of V1. Accordingly, all analyzed receptive fields were located below the horizontal meridian and contralateral to the recorded hemisphere: the left visual field for monkey B and the right visual field for monkey J. Their eccentricities ranged from approximately 1.5° to 8°, with most located between 2° and 3°. Receptive fields closer to fixation could overlap the fixation target and its allowable gaze window; those penetrations were not pursued. The stereoscope restricted the usable horizontal stimulus field to approximately 10° eccentricity. For each acute penetration, stimulus location and diameter were selected to cover the mapped receptive fields without extending beyond the usable display region. Experimental grating diameters ranged from 0.5° to 2.5° and generally increased with eccentricity.

##### Receptive field identification, eye and orientation tuning

Receptive fields were initially localized using manually positioned gratings and then assessed with localized, Gaussian-windowed binary-noise patches. The noise consisted of fine bright and dark pixels, rather than a sinusoidal grating with a defined spatial frequency. The mapping code sampled eccentricities from 2° to 6° in 1° increments and polar angles from 0° to 315° in 35° increments. Noise-patch diameter was scaled with eccentricity, and the code specified 200-ms presentations separated by 200-ms intervals. Grating tuning was then measured with stimuli shown binocularly or monocularly to either eye. For the dataset reported here, orientation and eye tuning were performed with drifting gratings rather than static gratings. The tuning script sampled 16 orientations (0°–168.75° in 11.25° increments); its nominal settings were 8-Hz drift, 2 cycles/degree, and 90% Michelson contrast. A separate spatial-frequency tuning routine offered 0.25, 0.5, 1, and 2 cycles/degree. Stimulus settings could be adjusted for each penetration according to receptive-field position and size.

##### Recording-site eligibility and offline preference classification

We used receptive-field overlap and alignment of preferred orientation along the putative V1 segment of each probe to evaluate whether the electrode sampled a single cortical column. Recordings proceeded only when more than 80% of V1 recording sites had aligned orientation preferences and overlapping receptive fields. This restriction was important because the 32-contact array extended approximately 3.1 mm between its most superficial and deepest contacts, whereas V1 can be as thin as approximately 1.5 mm. Deep contacts could therefore encounter other cortical folds with different receptive-field locations, orientation preferences, and eye preferences. Such sites were excluded; only contacts conservatively assigned to V1 were considered. The orientation preferred by the majority of eligible contacts was used to set the preferred and orthogonal orientations presented during the BRFS task. After acquisition, each recording site’s preferred eye and orientation were determined from its own monocular responses within the BRFS dataset. Trials were classified relative to that site’s preferences for analysis.

##### BRFS conditions, contrast, and trial inclusion

The main experimental stimuli were static sinusoidal gratings, usually at a spatial frequency of 1 cycle/degree and with 50% Michelson contrast. The contrast was equal in both eyes whenever gratings were simultaneously presented; this stimulus contrast was not intended to establish that V1 responses were below saturation. The experimental design included 20 conditions: four simultaneous binocular presentations, eight monocular-adaptation/binocular-flash presentations, and eight physical-alternation controls. Both preferred and orthogonal orientations were presented in balanced eye configurations.

MonkeyLogic generated pseudorandom condition sequences using draws with replacement; therefore, identical completed-trial counts were not guaranteed. Each BRFS trial contained 800 ms of monocular adaptation immediately followed by 800 ms of dichoptic stimulation, during which the adapted eye’s stimulus remained visible. In monocular replay, the first stimulus was extinguished as the new stimulus appeared in the other eye. Trials terminated upon fixation loss and were not counted as successfully completed. At least 30 successfully completed trials per condition were required for a session to be analyzed.

#### c. Data Preprocessing

For each recording session, linear multielectrode arrays (32 contacts, 0.1 mm apart; UProbe, Plexon Inc., Dallas, TX -or 32 contacts, 0.1 mm apart; Vector Array, NeuroNexus, Ann Arbor, MI) were implanted and removed. These arrays recorded extracellular voltage fluctuations, referenced against building ground. The Cerebus neural signal processing system (Blackrock Microsystems, Salt Lake City, UT), equipped with 128 channels, collected and processed the signals, amplifying and filtering the voltage time series for offline analysis. The raw signals, were sampled at 30 kHz, with a frequency range of 0.3-7.5 kHz.

The local field potential (LFP) was a down-sampled to 1 kHz from the raw broadband signal. Current Source Density (CSD) was calculated using an approximation of the second spatial derivative of broad-band LFP with the following formula (45, 46, 118):

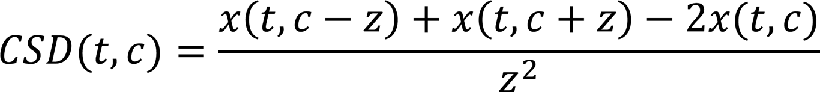

Where *x* is the extracellular voltage at time *t* measured at an electrode contact at position *c,* and *z* is the inter-contact distance of the electrode array. The resulting CSD from the formula above was multiplied by 0.4 S/mm as an estimate of the electric conductivity of cortex (119) to obtain the current per unit volume.

Power spectral density (PSD) was obtained separately for each penetration. PSD was calculated using a custom MATLAB script based for the fast Fourier transform (FFT). For each channel, we applied a Hanning window (n = 512 points, 50% overlap) to a time-segment of the local field potential (LFP) and computed the FFT, taking the squared magnitude of the resulting complex values to obtain the power spectrum. We retained only the positive frequency components (0-150Hz). Power spectra were computed for individual trials and then averaged across trials to obtain a mean power spectrum for each channel. To normalize across channels, we used the following formula (47):

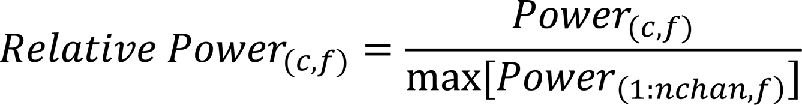

where *c* is each channel on the probe, and *f* is each frequency from 0Hz to 150Hz. For each frequency bin, the channel, with the maximum power was assigned a value of 1, and all other channels were scaled relative to that maximum. This results in a two-dimensional relative power matrix (channel x frequency) for each probe. The crisscross of the gamma-beta power spectrum is a reliable indicator of the cortical column’s granular input layer (47).

Coherence analysis was used to examine the synchronization between pairs of electrode contacts within V1. The *mscohere.m* function in MATLAB was employed to assess magnitude-squared coherence, providing insights into the dynamic interactions within the neural circuits. Coherence was evaluated on LFP data from the last 512ms trials because this time-window of sustained firing yields the most stationary response to avoid violating assumptions of the Fourier transform. The magnitude-squared coherence estimate *C_xy_(f)* indicates how well *x* corresponds to *y* on a scale of 0 to 1, and is obtained by:

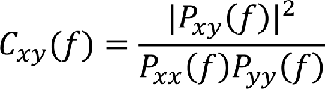

Where *P_xx_*(*f*)and *P_yy_*(*f*) are power spectral densities and *P_xy_*(*f*) is the cross power spectral density for each electrode pair *x* and *y*. We took the median coherence value from 1-30Hz on each trial for each electrode pair, and then averaged this coherence value across trials and recording penetrations.

To extract multi-unit activity (MUA) from the broadband signal, the voltage signal was band-pass filtered between 750 and 5000 Hz using a bidirectional Butterworth filter. The signal was then full-wave rectified and low-pass filtered at 500 Hz to yield the envelope of the raw voltage signal (50).

#### d. Laminar Analyses of Each Probe

We initially evaluated 44 acute laminar penetrations (12 from monkey B and 32 from monkey J). Laminar alignment was performed using established electrophysiological markers (26, 43, 45, 46). The bottom of granular layer IVc was identified using the initial stimulus-evoked CSD sink (45, 46) and corroborated by the transition between alpha–beta power in deeper cortex and theta–gamma power in upper cortex (47), as well as the characteristic discontinuity in interlaminar coherence (26, 43).

Receptive fields were mapped along each probe to verify their overlap, and the predominant orientation preference was evaluated across recording contacts. We proceeded with experimental recordings only when more than 80% of the putative V1 sites showed aligned orientation tuning and overlapping receptive fields, consistent with an approximately orthogonal penetration through a single cortical column (14, 43). This criterion was especially important because the 32-contact electrode spanned approximately 3.1 mm, whereas V1 can be as thin as approximately 1.5 mm; deeper contacts could therefore enter another cortical fold and exhibit markedly different receptive-field, orientation, or ocular preferences. We conservatively excluded contacts not confidently assigned to V1 and penetrations without reliable alignment of the V1 laminae. Based on the receptive-field, tuning, and laminar-alignment criteria, 31 of the original 44 penetrations were retained. We next required at least five consecutive electrode contacts, spaced 100 µm apart, in each operationally defined 0.5-mm compartment: supragranular, granular, and infragranular. These compartments approximately corresponded to layers II/III, IV, and V/VI, respectively, and were defined physiologically rather than by histology. Four of the 31 aligned penetrations did not span all three compartments sufficiently and were therefore excluded from the compartment-based analyses. The final dataset comprised 27 penetrations (eight from monkey B and 19 from monkey J), with 15 selected V1 contacts per penetration and 405 MUA recording sites in total (135 per compartment).

## Supplemental figures

**Supplemental Figure 1.**
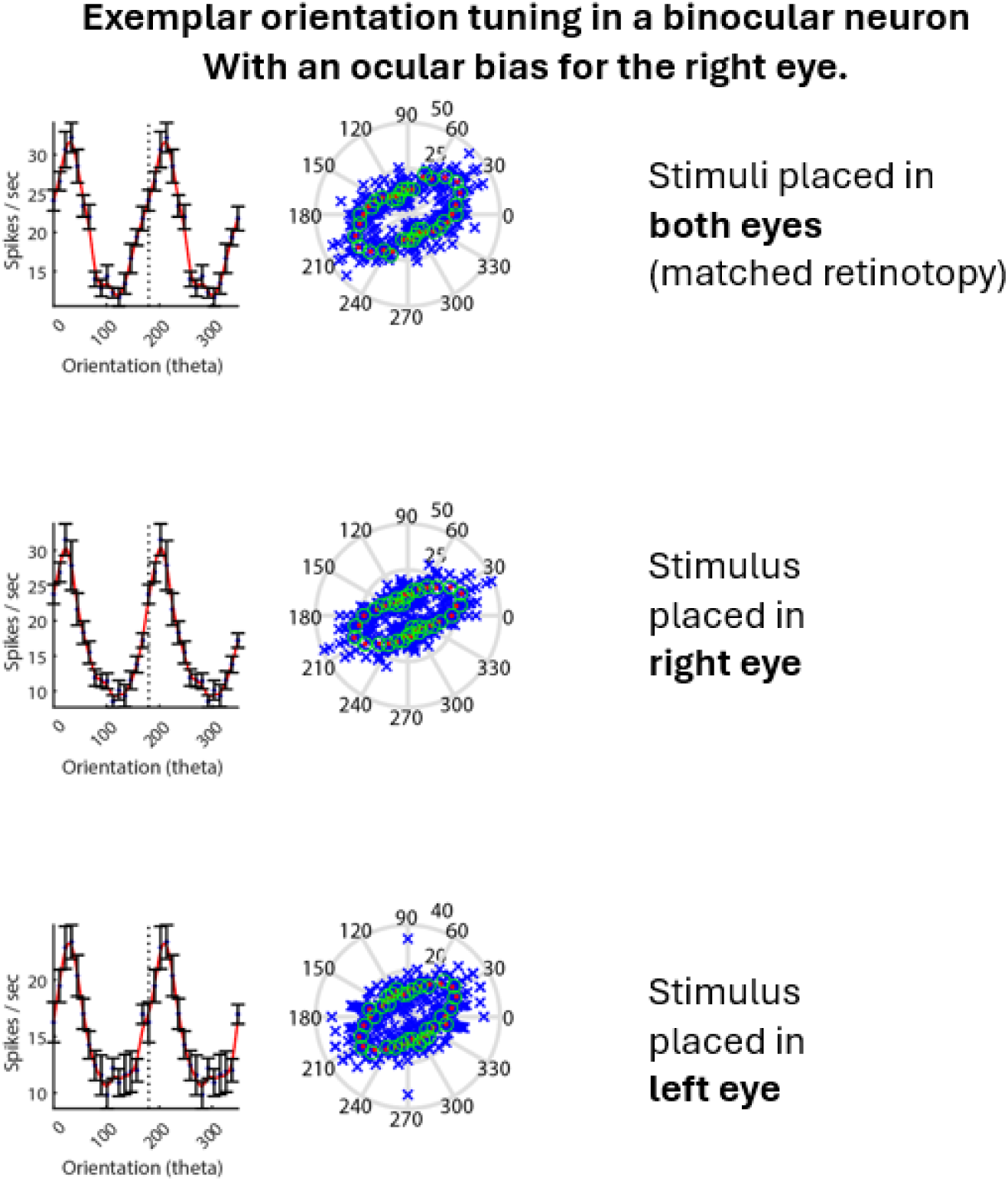
Example orientation and ocular tuning of a V1 multiunit. Following identification of laminar boundaries and isolation of multiunit activity, we assessed orientation and ocular preferences by presenting gratings of varying orientations monocularly to each eye. This example recording site demonstrates a preference for stimulation of the right eye and an orientation preference of approximately 30°.

**Supplemental Figure 2.**
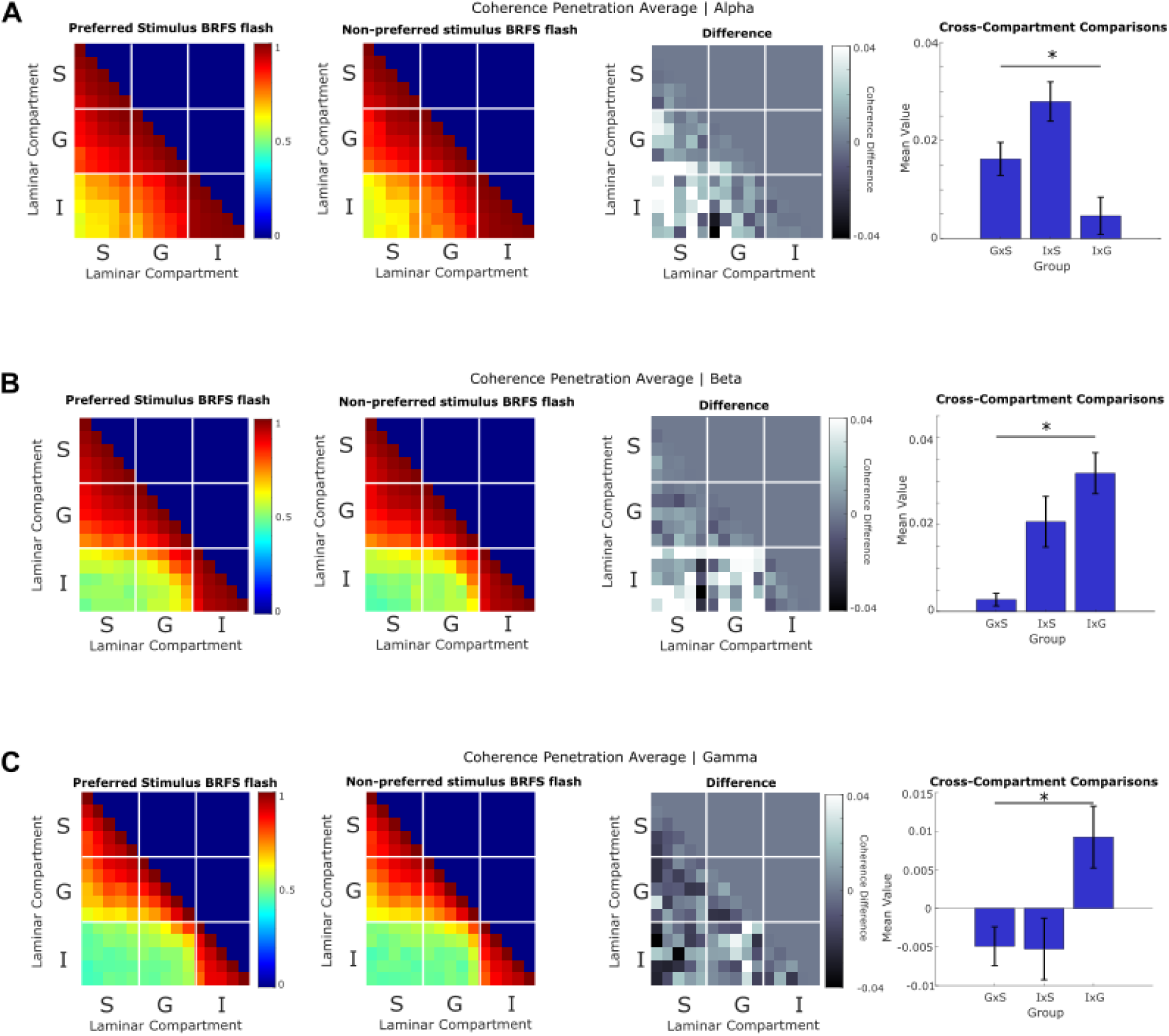
BRFS interlaminar coherence across different frequency bands. Magnitude-squared coherence was calculated during the last 512 ms of binocular rivalry flash suppression (BRFS) for three frequency bands: (A) alpha (7.8–11.7 Hz), (B) beta (15.6–27.3 Hz), and (C) gamma (31.3–58.6 Hz). Each row shows the mean interlaminar coherence for BRFS when the preferred stimulus was presented second, when the null stimulus was presented second, and the difference between these conditions. The rightmost plots show the distribution of coherence differences across the three interlaminar compartment comparisons. The alpha-and beta-band analyses reveal patterns of interlaminar coherence differences similar to those reported in Figure 4, which evaluated coherence over 1–30 Hz. The gamma-band analysis also reveals a significant effect of interlaminar compartment in the omnibus ANOVA, although the coherence differences are in the opposite direction from those observed at lower frequencies. The asterisk indicates a significant effect of interlaminar compartment on the coherence differences between conditions (one-way ANOVAs; alpha: *F*(2,72) = 9.70, p < 0.001; beta: *F*(2,72) = 11.13, p < 0.001; gamma: *F*(2,72) = 5.35, p = 0.007).

**Supplemental Figure 3.**
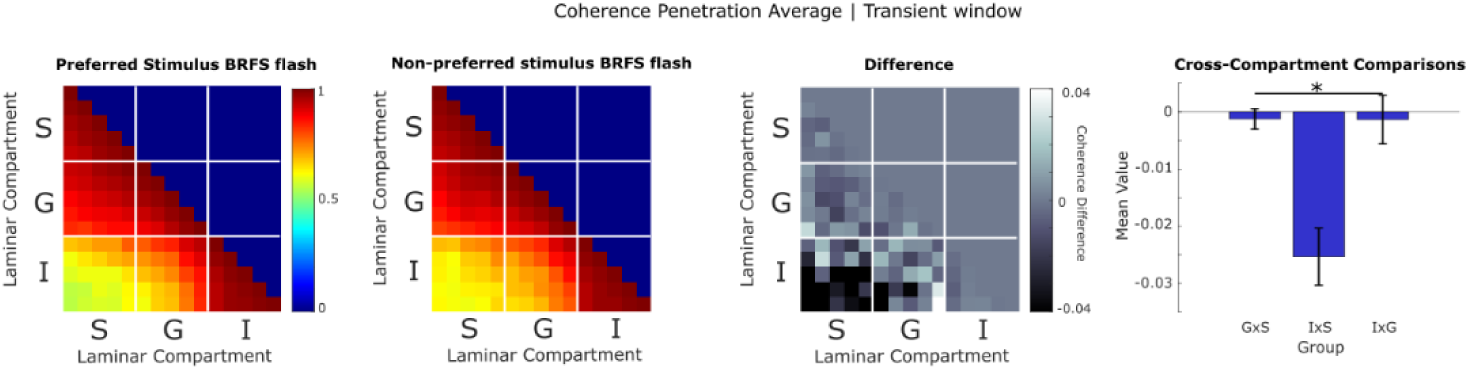
BRFS interlaminar coherence during the early stimulus-onset transient. Magnitude-squared coherence was calculated over the first 512 ms following the onset of dichoptic stimulation during BRFS. As in Figure 4, the coherence matrices show when the preferred stimulus was flashed, when the null stimulus was flashed, and the difference between these conditions. The rightmost plot shows the distribution of coherence differences across the three interlaminar compartment comparisons. Note that the negative mean values for interlaminar coherence indicate that, within the transient window, the non-preferred BRFS flash has more coherence than the preferred BRFS flash. This relationship is the opposite of what was observed during the sustained time-window. The asterisk indicates a significant effect of interlaminar compartment on the coherence differences between conditions (one-way ANOVA, *F*(2,72) = 12.44, p < 0.001) Unlike the sustained response window used in the primary analyses, this interval includes the large transient neuronal response evoked by the dichoptic flash. These results should be interpreted cautiously because stimulus-evoked transients introduce nonstationarity that can influence Fourier-based estimates of coherence.

**Supplemental Figure 4.**
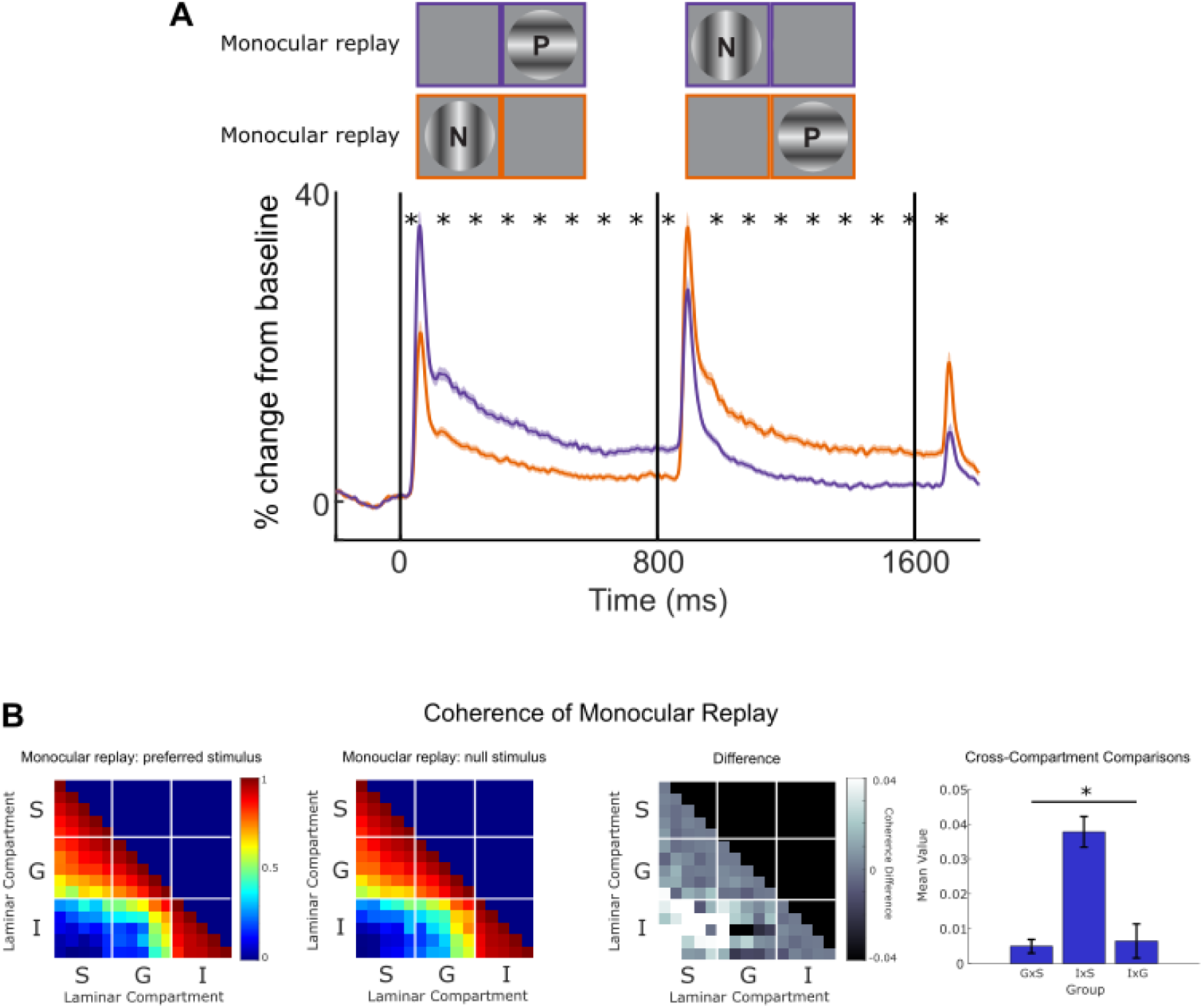
Population multiunit activity during monocular replay. (A) Population multiunit activity (MUA) during the monocular replay control condition, which physically alternated stimuli between the eyes to reproduce the expected perceptual sequence of BRFS without dichoptic stimulation. The cartoon above the MUA traces depicts the monocular stimulus sequence presented to the subject. The purple trace shows the preferred stimulus (eye and orientation) presented first, followed by the null stimulus. The orange trace shows the opposite sequence, with the null stimulus presented first, followed by the preferred stimulus. Solid lines represent the mean MUA response, expressed as a percent change from baseline. Asterisks indicate 100-ms time bins with a statistically significant difference between conditions (one-tailed paired t-tests, Bonferroni-corrected for 16 comparisons; α = 0.05). (B) Magnitude-squared coherence was calculated during the last 512 ms of visual stimulation. Coherence matrices show the mean interlaminar coherence for the two compared conditions, their difference, and the distribution of coherence differences across the three interlaminar compartment comparisons. Interlaminar coherence during monocular replay. The asterisk indicates a significant effect of interlaminar compartment on the coherence differences between conditions (one-way ANOVA, *F*(2,72) = 21.93, p < 0.001), similar to the pattern observed during BRFS in Figure 4. However, average coherence between the granular and infragranular layers was lower during monocular replay than during BRFS.

**Supplemental Figure 5.**
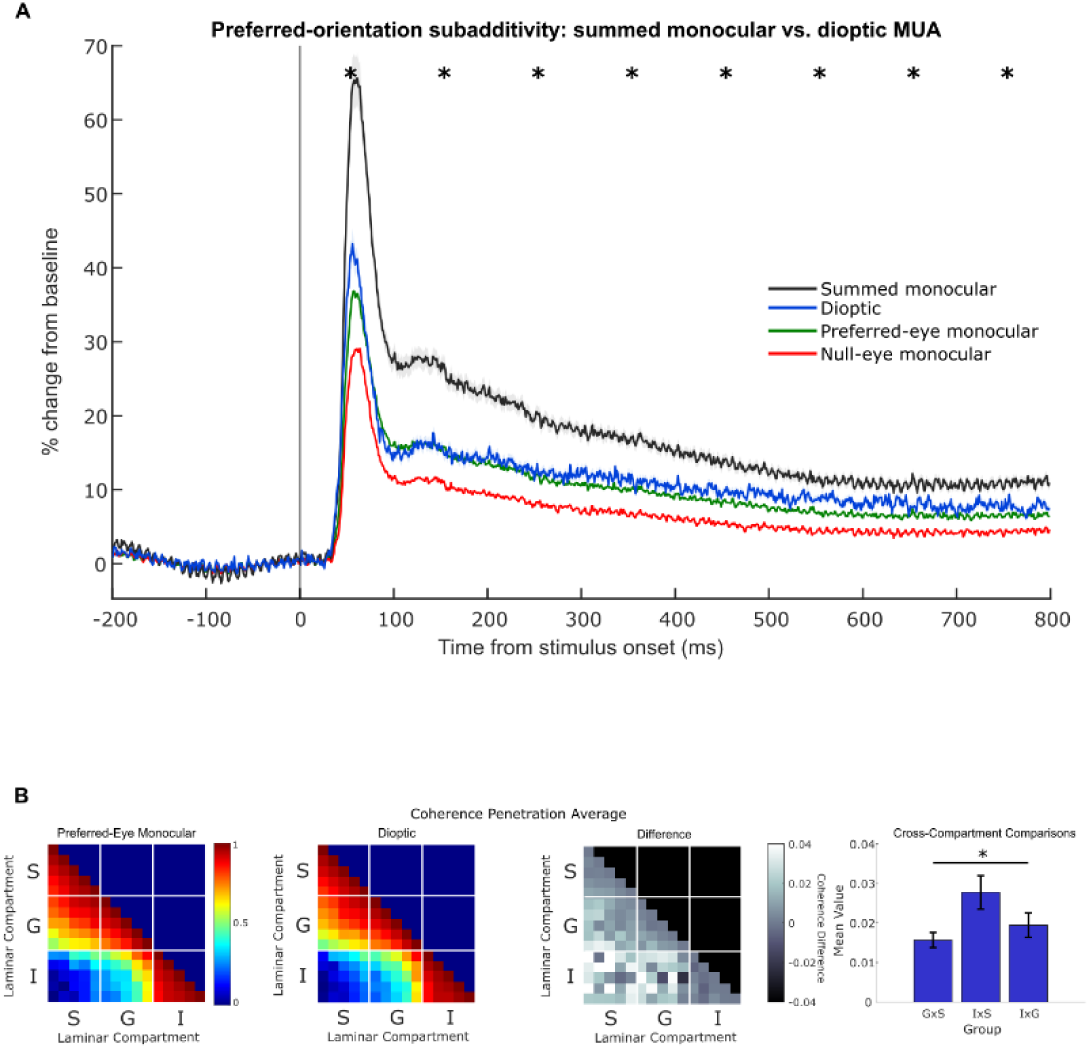
Subadditive neuronal responses during binocular stimulation. (A) Grand-average multiunit activity (MUA) evoked by monocular stimulation of the preferred eye (green), monocular stimulation of the non-preferred eye (red), and simultaneous dioptic stimulation of both eyes (blue). All conditions presented the preferred stimulus orientation. The black trace represents the linear sum of the two monocular responses, illustrating the predicted binocular response in the absence of subadditive interactions. Responses are plotted as mean percent change from baseline, with shaded regions indicating the standard error of the mean (SEM) for the summed monocular and dioptic responses. Asterisks indicate 100-ms time bins in which the summed monocular response was significantly greater than the dioptic response (one-tailed paired t-tests, Bonferroni-corrected for 8 comparisons; α = 0.05). All eight time-bins showed significant subadditivity. The dioptic response was substantially smaller than the linear sum of the monocular responses and closely resembled the preferred-eye monocular response, consistent with binocular normalization. An additional paired, two-tailed t-test revealed that preferred-eye monocular responses were significantly lower than dioptic responses during the sustained 512-ms window (288–800 ms; mean difference = −1.76 percentage points, t(404) = −6.44, p < 0.001). Bonferroni-corrected comparisons across eight 100-ms time bins revealed significant differences in seven bins (α = 0.00625). (B) Interlaminar coherence during preferred-eye monocular and dioptic stimulation. Magnitude-squared coherence was calculated during the final 512 ms of the initial stimulus presentation (288–800 ms). Coherence matrices show the mean interlaminar coherence during monocular and dioptic stimulation, followed by their difference (monocular minus dioptic). The rightmost plot shows coherence differences across the three interlaminar compartment comparisons. Coherence differences varied significantly across compartments (one-way ANOVA, F(2, 72) = 3.64, p = 0.031). The asterisk indicates a significant main effect of interlaminar compartment (α = 0.05).

## ACKNOWLEDGMENTS

The authors would like to thank Randolph Blake, PhD and Catie Chang, PhD, for their comments on an earlier version of this work.

“Preprint is available at https://doi.org/10.1101/2025.10.04.680461”

## AUTHOR CONTRIBUTIONS

B.M.C and A.M designed research; B.M.C., B.A.M, and J.A.W performed experiments; B.M.C., B.A.M, and P.G.P analyzed data; B.M.C., B.A.M, J.A.W., G.W, and A.M. interpreted results of the experiments; B.M.C. prepared figures; B.M.C. drafted manuscript; B.M.C., B.A.M., J.A.W., P.G.P., G.W., and A.M edited and revised manuscript; B.M.C., B.A.M., J.A.W., P.G.P., G.W., and A.M approved final version of manuscript.

