## Supplemental figures for "V1 interlaminar coherence decreases with interocular conflict"

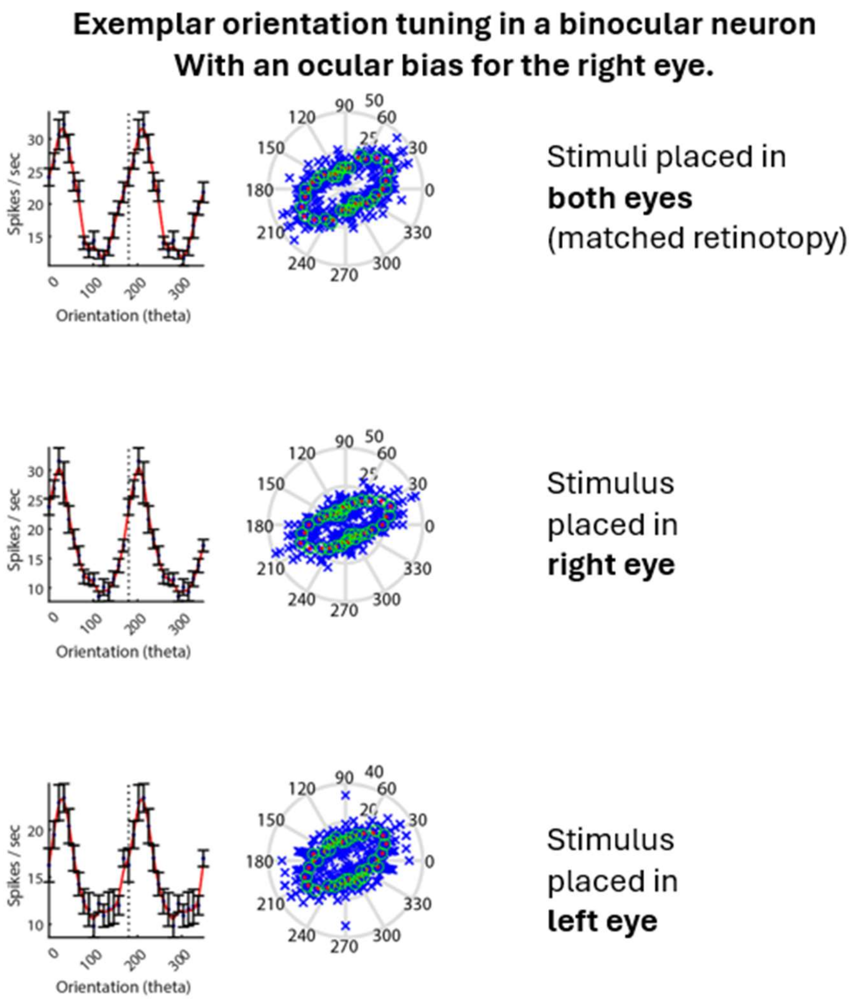


**Supplemental Figure 1.** Example orientation and ocular tuning of a V1 multiunit. Following identification of laminar boundaries and isolation of multiunit activity, we assessed orientation and ocular preferences by presenting gratings of varying orientations monocularly to each eye. This example recording site demonstrates a preference for stimulation of the right eye and an orientation preference of approximately 30°.

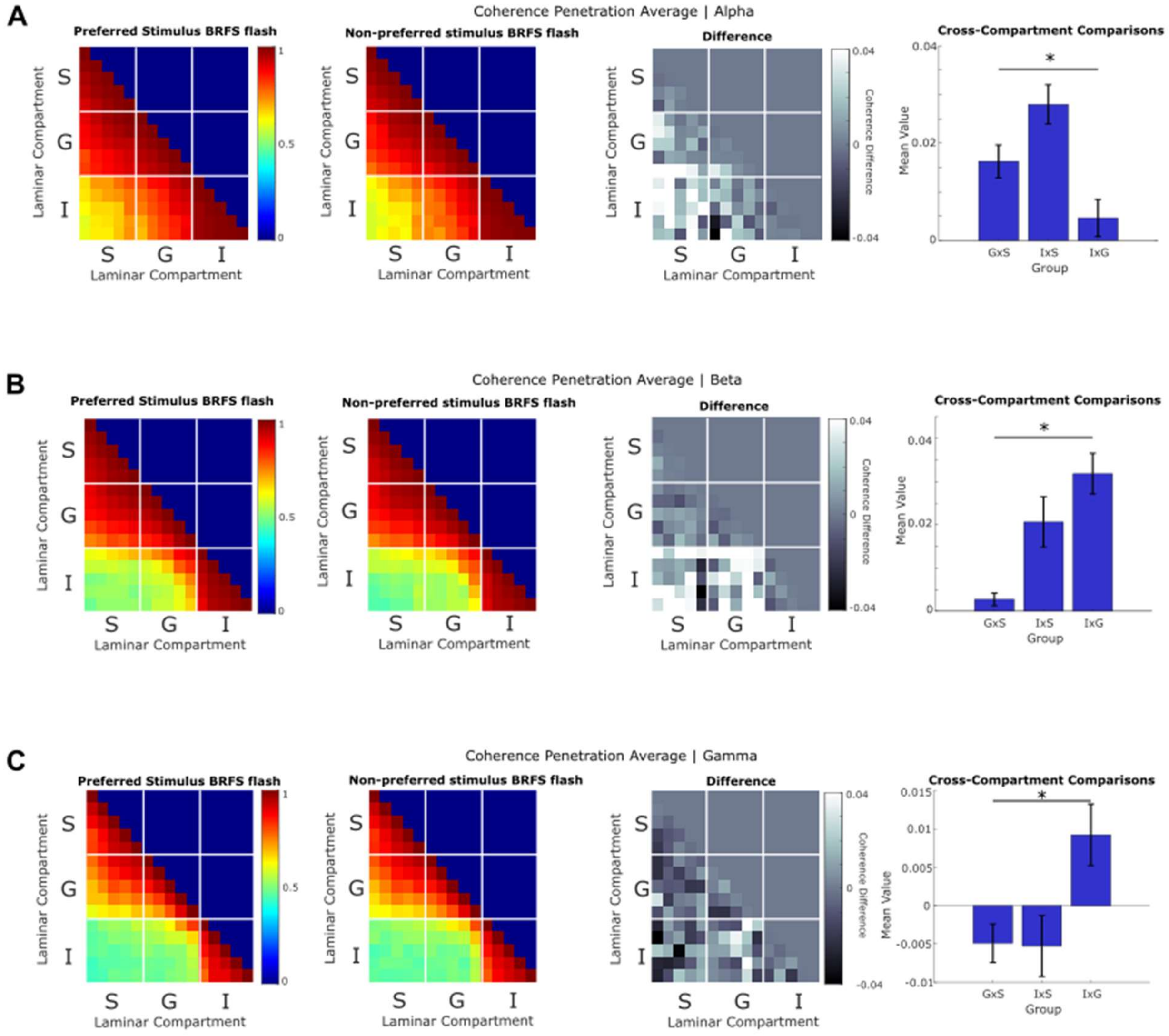


**Supplemental Figure 2. BRFS interlaminar coherence across different frequency bands.** Magnitude-squared coherence was calculated during the last 512 ms of binocular rivalry flash suppression (BRFS) for three frequency bands: (A) alpha (7.8–11.7 Hz), (B) beta (15.6–27.3 Hz), and (C) gamma (31.3–58.6 Hz). Each row shows the mean interlaminar coherence for BRFS when the preferred stimulus was presented second, when the null stimulus was presented second, and the difference between these conditions. The rightmost plots show the distribution of coherence differences across the three interlaminar compartment comparisons. The alpha- and beta-band analyses reveal patterns of interlaminar coherence differences similar to those reported in Figure 4, which evaluated coherence over 1–30 Hz. The gamma-band analysis also reveals a significant effect of interlaminar compartment in the omnibus ANOVA, although the coherence differences are in the opposite direction from those observed at lower frequencies. The asterisk indicates a significant effect of interlaminar compartment on the coherence differences between conditions (one-way ANOVAs; alpha: $F(2,72) = 9.70$, $p < 0.001$; beta: $F(2,72) = 11.13$, $p < 0.001$; gamma: $F(2,72) = 5.35$, $p = 0.007$).

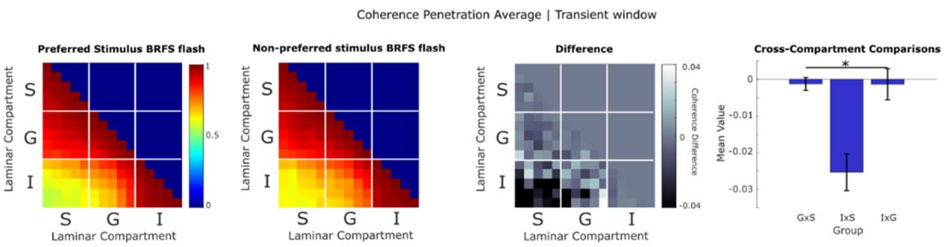


**Supplemental Figure 3. BRFS interlaminar coherence during the early stimulus-onset transient.** Magnitude-squared coherence was calculated over the first 512 ms following the onset of dichoptic stimulation during BRFS. As in Figure 4, the coherence matrices show when the preferred stimulus was flashed, when the null stimulus was flashed, and the difference between these conditions. The rightmost plot shows the distribution of coherence differences across the three interlaminar compartment comparisons. Note that the negative mean values for interlaminar coherence indicate that, within the transient window, the non-preferred BRFS flash has more coherence than the preferred BRFS flash. This relationship is the opposite of what was observed during the sustained time-window. The asterisk indicates a significant effect of interlaminar compartment on the coherence differences between conditions (one-way ANOVA, $F(2,72) = 12.44$, $p < 0.001$) Unlike the sustained response window used in the primary analyses, this interval includes the large transient neuronal response evoked by the dichoptic flash. These results should be interpreted cautiously because stimulus-evoked transients introduce nonstationarity that can influence Fourier-based estimates of coherence.

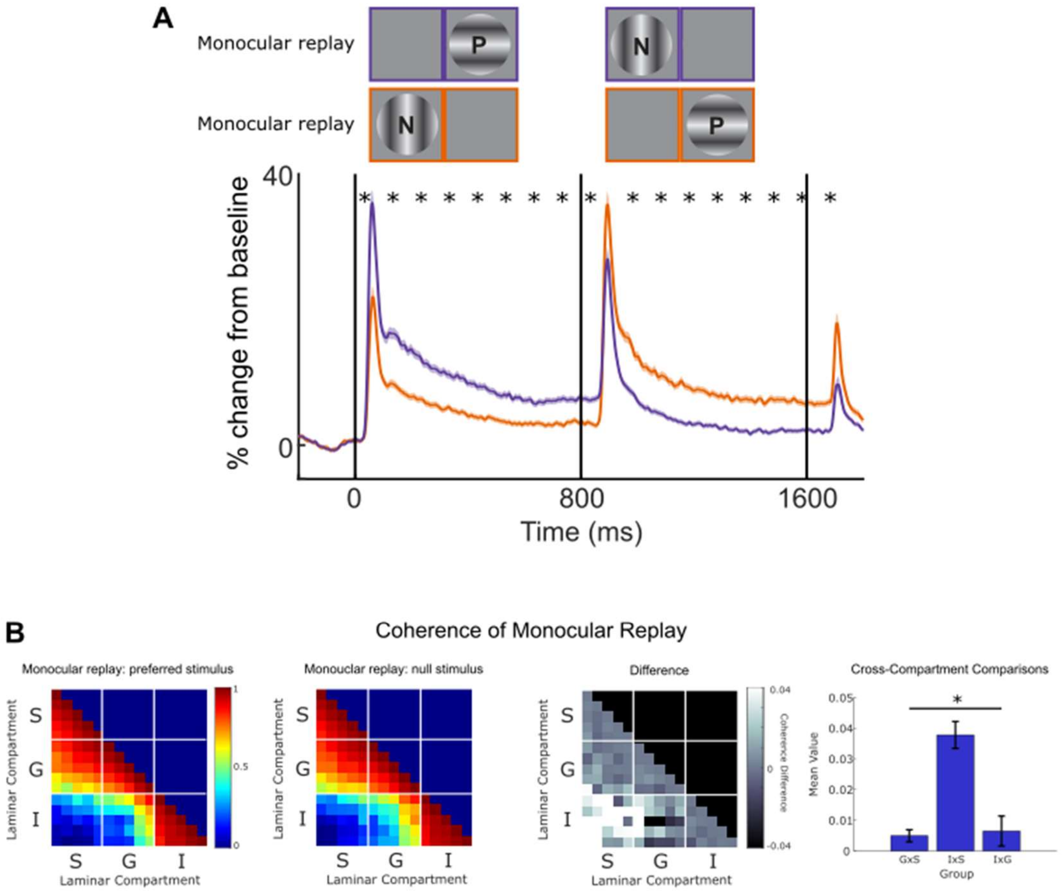


**Supplemental Figure 4. Population multiunit activity during monocular replay.** (A) Population multiunit activity (MUA) during the monocular replay control condition, which physically alternated stimuli between the eyes to reproduce the expected perceptual sequence of BRFS without dichoptic stimulation. The cartoon above the MUA traces depicts the monocular stimulus sequence presented to the subject. The purple trace shows the preferred stimulus (eye and orientation) presented first, followed by the null stimulus. The orange trace shows the opposite sequence, with the null stimulus presented first, followed by the preferred stimulus. Solid lines represent the mean MUA response, expressed as a percent change from baseline. Asterisks indicate 100-ms time bins with a statistically significant difference between conditions (one-tailed paired t-tests, Bonferroni-corrected for 16 comparisons; $\alpha = 0.05$). (B) Magnitude-squared coherence was calculated during the last 512 ms of visual stimulation. Coherence matrices show the mean interlaminar coherence for the two compared conditions, their difference, and the distribution of coherence differences across the three interlaminar compartment comparisons. Interlaminar coherence during monocular replay. The asterisk indicates a significant effect of interlaminar compartment on the coherence differences between conditions (one-way ANOVA, $F(2,72) = 21.93$, $p < 0.001$), similar to the pattern observed during BRFS in Figure 4. However, average coherence between the granular and infragranular layers was lower during monocular replay than during BRFS.

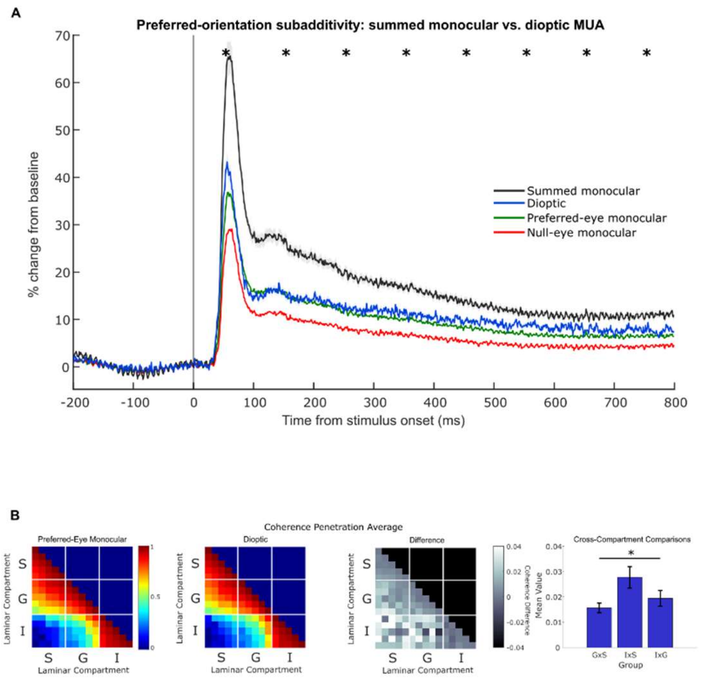


**Supplemental Figure 5. Subadditive neuronal responses during binocular stimulation.** (A) Grand-average multiunit activity (MUA) evoked by monocular stimulation of the preferred eye (green), monocular stimulation of the non-preferred eye (red), and simultaneous dioptic stimulation of both eyes (blue). All conditions presented the preferred stimulus orientation. The black trace represents the linear sum of the two monocular responses, illustrating the predicted binocular response in the absence of subadditive interactions. Responses are plotted as mean percent change from baseline, with shaded regions indicating the standard error of the mean (SEM) for the summed monocular and dioptic responses. Asterisks indicate 100-ms time bins in which the summed monocular response was significantly greater than the dioptic response (one-tailed paired t-tests, Bonferroni-corrected for 8 comparisons; $\alpha = 0.05$). All eight time-bins showed significant subadditivity. The dioptic response was substantially smaller than the linear sum of the monocular responses and closely resembled the preferred-eye monocular response, consistent with binocular normalization. An additional paired, two-tailed t-test revealed that preferred-eye monocular responses were significantly lower than dioptic responses during the sustained 512-ms window (288–800 ms; mean difference = −1.76 percentage points, $t(404) = -6.44$, $p < 0.001$). Bonferroni-corrected comparisons across eight 100-ms time bins revealed significant differences in seven bins ($\alpha = 0.00625$). (B) Interlaminar coherence during preferred-eye monocular and dioptic stimulation. Magnitude-squared coherence was calculated during the final 512 ms of the initial stimulus presentation (288–800 ms). Coherence matrices show the mean interlaminar coherence during monocular and dioptic stimulation, followed by their difference (monocular minus dioptic). The rightmost plot shows coherence differences across the three interlaminar compartment comparisons. Coherence differences varied significantly across compartments (one-way ANOVA, $F(2, 72) = 3.64$, $p = 0.031$). The asterisk indicates a significant main effect of interlaminar compartment ($\alpha = 0.05$).
